# Morphological and sensorimotor phenotypes in a zebrafish CHARGE syndrome model are domain-dependent

**DOI:** 10.1101/2022.07.14.499979

**Authors:** Dana R. Hodorovich, Patrick M. Lindsley, Austen A. Berry, Derek F. Burton, Kurt C. Marsden

## Abstract

CHARGE syndrome is a rare disorder characterized by a spectrum of defects affecting multiple tissues and behavioral difficulties such as autism, attention-deficit/hyperactivity disorder, obsessive-compulsive disorder, anxiety, and sensory deficits. Most CHARGE cases arise from *de novo*, loss-of-function mutations in a master transcriptional regulator, chromodomain-helicase-DNA-binding-protein-7 (CHD7). CHD7 regulates key neurodevelopmental factors and is required for neural processes including neuronal differentiation and neural crest cell migration, but how CHD7 affects neural circuit function to regulate behavior is unclear. To investigate the pathophysiology of behavioral symptoms in CHARGE, we established a mutant *chd7* zebrafish line using CRISPR/Cas9 that recapitulates multiple CHARGE phenotypes. Using a panel of behavioral assays, we find that *chd7* mutants have specific auditory and visually driven behavioral deficits that are independent of defects in sensory structures, implicating *chd7* in the regulation of underlying brain circuits. Furthermore, by analyzing multiple *chd7* alleles we show that the penetrance of morphological and behavioral phenotypes depends on the mutation location. These results provide novel insight into the heterogeneity of CHARGE syndrome and will inform future work to define mechanisms of CHD7-dependent neurobehavioral symptoms.

**Summary statement:** Mutation location within *chd7* affects the expression of CHARGE Syndrome-related morphological and behavioral phenotypes in a larval zebrafish CHARGE model.

## Introduction

CHARGE syndrome is an autosomal dominant disorder that affects approximately 1 in every 10,000 births (Zentner et al., 2010a). CHARGE is an acronym describing the previously used diagnostic criteria, but disease diagnosis now includes the presentation of at least one major feature including coloboma of the eye, atresia of the choanae, and semicircular canals defects, or a major sign with additional minor symptoms including heart or esophageal anomalies (Verloes, 2005) cleft lip/palate, cranial nerve(s) dysfunction (Blake et al., 2008), and developmental delay (Blake and Prasad, 2006). In addition to these symptoms, ∼60% of CHARGE patients are prescribed medication for mood and/or behavioral difficulties (Blake et al., 2005). These include hypo- or hyper-sensory processing problems (Hartshorne et al., 2017), autistic-like behaviors (Hartshorne et al., 2005), Obsessive-compulsive disorder and repetitive behaviors (Bernstein and Denno, 2005), attention-deficit/ hyperactivity disorder (AD/HD) (Hartshorne et al., 2016), anxiety (Hartshorne et al., 2017), aggression (Blake et al., 2005), diminished self-regulation (Hartshorne et al., 2017), self-abusive behaviors (Blake et al., 2005) (Souriau et al., 2005), sleep issues (Hartshorne et al., 2016) and intellectual disability (Wachtel et al., 2007). Despite their frequency, the pathophysiology of behavioral symptoms in CHARGE patients is not well understood.

*De novo* loss-of-function mutations in the chromatin remodeling gene *Chromodomain-helicase-DNA-binding-protein 7* (CHD7) cause ∼70% of CHARGE cases (Vissers et al., 2004; Zentner et al., 2010b). Mutations have been found throughout the gene and include a variety of missense, nonsense, frameshift, and splice site mutations (Bergman et al., 2011; Jongmans et al., 2006; Legendre et al., 2017), indicating there is no hotspot for disease pathogenesis. CHD7 is an ATP-dependent transcriptional regulator ubiquitously expressed with enhanced expression in the developing brain (Sanlaville et al., 2006). CHD7 has multiple roles in neurodevelopment including neural crest cell migration and differentiation (Schulz et al., 2014; Wysocka et al., 2010), cerebellar organization and foliation (Whittaker et al., 2017a; Whittaker et al., 2017b), olfactory development (Bergman et al., 2010; Layman et al., 2009), regulation of myelination and remyelination factors (He et al., 2016), and glial and neuronal differentiation (Jones et al., 2015). Animal models of CHARGE syndrome have provided insight into the etiology of ear and vestibulocochlear defects (Bosman et al., 2005), gonadal deficits, interventricular heart deficits (Bergman et al., 2010; Bosman et al., 2005), gut motility dysfunction (Cloney et al., 2018), and craniofacial defects (Asad and Sachidanandan, 2020; Asad et al., 2016; Balow et al., 2013). Here we sought to build on these efforts, using CRISPR/Cas9 to generate a new zebrafish CHARGE model and define how CHD7 regulates sensorimotor behavior.

Our model recapitulates several CHARGE phenotypes including ear, heart, and craniofacial defects, and our data also show that *chd7* is required for specific visual and auditory behaviors. Loss of *chd7* alters behavioral selection following acoustic stimulation, a phenotype that is both context-dependent and independent of defects in sensory structures. Finally, by analyzing multiple zebrafish CHARGE models, we find that the location of the mutation within the *chd7* gene affects the penetrance of CHARGE-related morphological and behavioral deficits. Our data support the highly conserved functions of CHD7 and reflect the heterogeneity of CHARGE syndrome patients, provide new insight into the variation in disease presentation.

## Results

### CRISPR/Cas9-generated *chd7* mutants recapitulate CHARGE phenotypes

Zebrafish larvae are an excellent animal model for examining the mechanisms underlying developmental and behavioral deficits in CHARGE syndrome because of their shared genetics, including 82% of disease-related genes (Howe et al., 2013) and ∼68% amino acid identity for CHD7, accessibility to genetic manipulation, conserved vertebrate structures within the central nervous system, along with their small size and extensive behavioral repertoire that allows for high-throughput testing (Friedrich et al., 2010; Kalueff et al., 2013). Larval zebrafish CHARGE syndrome models have previously been generated through morpholino knock-down and more recently using CRISPR/Cas9, and have been used to investigate specific phenotypes such as craniofacial (Asad et al., 2016), heart (Patten et al., 2012), and gut defects (Cloney et al., 2018). To generate a stable *chd7* mutant line for investigating disease-related phenotypes and a panel of sensorimotor behaviors, we designed a single guide RNA (gRNA) targeting a highly conserved region in Exon 9. We microinjected this gRNA along with Cas9 protein into 1-cell stage TLF-strain zebrafish embryos. We raised injected embryos to adulthood, crossed them to wild-type TLF fish, and PCR-amplified and sequenced the gRNA target site in F1 offspring to assess germ-line transmission of CRISPR/Cas9-induced mutations. We identified a 7-base pair (bp) deletion that produces a frameshift and premature stop codon within the second chromodomain of Chd7 (Fig. 1A). First, we assessed the viability of *chd7* mutants until 30 days post fertilization (dpf) and found that by 15 dpf in contrast to the 90% survival rate of wildtypes, only 24% of mutants and 59% of heterozygotes survived (Fig. 1B), with no changes in viability thereafter. These data are similar to those showing embryonic lethality of homozygous *chd7* mutations in mammals (Bosman et al., 2005), but the semi-viability of our zebrafish *chd7* mutants establishes an opportunity to evaluate *chd7* function in a homozygous context. We next used RT-qPCR to measure *chd7* transcript levels in pools of 5 dpf larvae and observed that *chd7* mRNA was significantly downregulated by 22% in heterozygotes and by 54% in mutants, verifying the deleterious impact of the mutation, with the residual transcript likely reflecting maternal deposition and/or incomplete nonsense-mediated decay (Fig. 1C).

**Figure 1.**
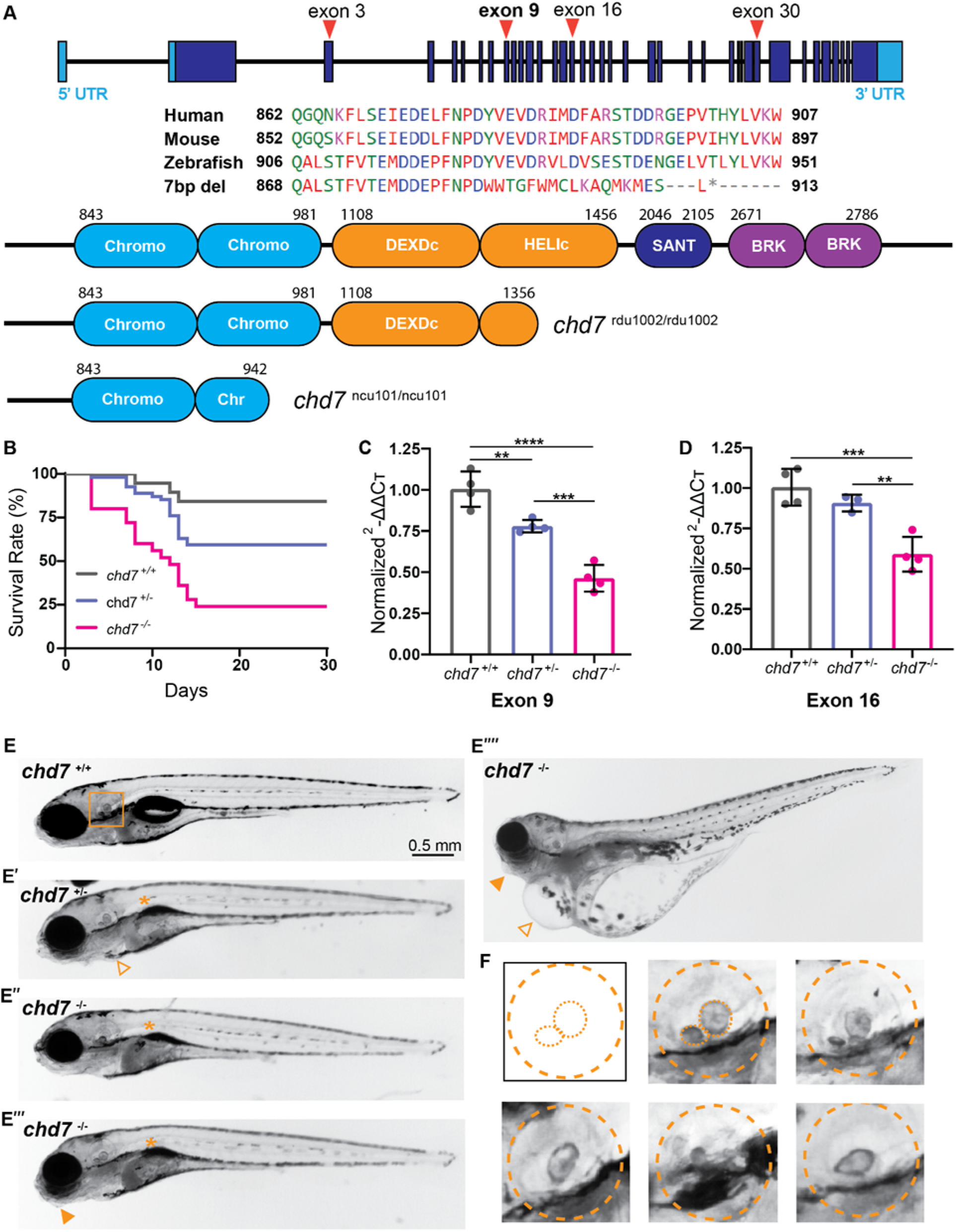
CRISPR/Cas9 generated *chd7* mutants recapitulate CHARGE syndrome related phenotypes. **(A)** Exon-intron structure of zebrafish *chd7* with CRISPR targeted regions noted; amino acid alignment and conservation of Exon 9 target across species, identified 7 base pair (bp) deletion from mutagenesis; predicted chd7 protein domains in wild-type (top), Exon 16, 1 bp deletion allele (*chd7* ^*rdu1002/rdu1002*^), and Exon 9, 7 bp deletion allele (*chd7* ^*ncu101/ncu101*^), additional target regions from sgRNAs noted. **(B)** Survival rate (%) of *chd7*^+/+^ (n=19), *chd7*^+/-^ (n=54), and *chd7*^-/-^ (n=25) siblings during a 30-day period. **(C)** qPCR analysis of *chd7* mRNA in 5 dpf larvae (*chd7* ^*ncu101/*^, Exon 9 allele), normalized to wild-type siblings. Each point represents a biological replicate of pooled larvae (per pool, *chd7*^*+/+*^ n∼12-13; *chd7*^*+/-*^ n∼20-21; *chd7*^*-/-*^ n∼13-14). **(D)** qPCR analysis of *chd7* mRNA in 5 dpf larvae (*chd7* ^*rdu1002/*^, Exon 16 allele), normalized to wild-type siblings. Each point represents a biological replicate of pooled larvae (per pool, *chd7*^*+/+*^ n∼13-14; *chd7*^*+/-*^ n∼22-23; *chd7*^*-/-*^ n∼11-12) (mean ± SD, Ordinary one-way ANOVA with Tukey’s multiple comparisons, *p<0.05, **p<0.01, ***p<0.001,****p<0.0001,).

To determine whether our larval zebrafish model recapitulates CHARGE-related developmental phenotypes, we examined the morphology of 5 dpf *chd7* larvae. Both mutants and heterozygous siblings frequently displayed craniofacial defects, uninflated swim bladders, enlarged pericardial area, and ear defects (Fig. 1E). We did not observe differences in body length or eye size. In teleost fish the otoliths, hard crystal-like structures within the otic vesicle, are tethered to the sensory hair cells and function in hearing and balance (Nicolson, 2005; Schulz-Mirbach et al., 2019). We observed a variety of otolith phenotypes in *chd7* mutants and heterozygotes, including small anterior otoliths, missing anterior or posterior otoliths, and a merged or misshapen phenotype (Fig. 1E). These defects occurred both unilaterally and bilaterally. Otolith defects were the most penetrant phenotype, with 29% of heterozygotes and 65% of mutants displaying a phenotype. This reflects the case with human CHARGE patients, as inner and outer ear defects are the most common phenotype with at least 80% of patients reporting ear anomalies across clinical studies (Blake and Prasad, 2006). Overall, 45.1% of heterozygotes and 87.5% of mutants displayed at least one morphological phenotype (Table 1).

**Table 1.** Morphological phenotype frequencies in Exon 9 mutants. Morphological phenotype ratio and frequencies in 5-6 dpf *chd7*^*ncu101/+*^ *and chd7*^*ncu101/ncu101*^.

|  | Craniofacial defect | Otolith defect | Pericardial edema | Noninflated swim bladder | Larvae with any morphological phenotype |
| --- | --- | --- | --- | --- | --- |
| *chd7*<sup>+/-</sup> | 14/71, 19.7% | 20/71, 28.8% | 7/71, 9.9% | 4/71, 5.6% | 32/71, 45.1% |
| *chd7*<sup>-/-</sup> | 17/40, 42.5% | 26/40, 65% | 15/40, 37.5% | 18/40, 45% | 35/40, 87.5% |

To verify the impact of germline mutations in *chd7* on these developmental phenotypes, we analyzed an independent *chd7* mutant line that was generated in the AB wild-type strain using CRISPR/Cas9 with a gRNA targeting Exon 16 (Fig. 1A; gifted by Dr. Erica Davis, Northwestern University). The identified 1 bp deletion creates a premature stop codon within the ATP-helicase domain. We observed similar morphological phenotypes in Exon 16 mutants, but with decreased penetrance in heterozygotes. In contrast to Exon 9 larvae, there was a striking absence of otolith defects in both Exon 16 heterozygous and mutant larvae (Table S1). To test possible explanations for the difference in otolith phenotypes between the two mutant lines, we investigated whether there were CRISPR-mediated off-target mutations of the Exon 9 gRNA. DNA from F_1_ and F_2_ *chd7* heterozygous adults was assessed by PCR and Sanger sequencing for the top three predicted off-target locations (via chopchop.cbu.uib.no). There were no alterations to the base sequence of these potential off-target sites (Fig. S1A). Finally, using RT-qPCR we found that *chd7* transcript levels in Exon 16 mutants were significantly downregulated (Fig. 1D), as in Exon 9 mutants. Since both the Exon 9 and Exon 16 mutants survive through larval stages, have decreased *chd7* transcript, and recapitulate multiple but incompletely overlapping sets of disease-related phenotypes, these mutant lines provide a useful set of tools to study how different *chd7* mutations impact development and behavior.

### *chd7* is required for the central processing of auditory stimuli in the LLC circuit

Clinical studies have reported that at least 80% of CHARGE patients present with an external ear defect and either conductive or sensorineural hearing loss (Blake and Prasad, 2006). To determine if auditory detection was intact in our zebrafish CHARGE model, we performed an acoustic startle assay in 5 dpf larvae generated by crossing Exon 9 heterozygous adults. Larvae received 60 acoustic stimuli, 10 trials at each of 6 intensities, with a 20 second interstimulus interval (ISI). Following an acoustic stimulus, zebrafish larvae perform one of two response types: 1) Short-latency c-bends (SLCs), which are initiated 4-15 ms after the stimulus and are driven by a pair of command-like reticulospinal neurons known as the Mauthner cells (Burgess and Granato, 2007a), and 2) Long-latency c-bends (LLCs), which are initiated at least 15 ms after the stimulus, have reduced angular velocity, are independent of the Mauthner cells, and are mediated by a cluster of prepontine neurons in rhombomere 1 (Burgess and Granato, 2007a; Marquart et al., 2019). Larval zebrafish bias their responses toward SLCs following strong acoustic stimuli and toward LLCs following weaker stimuli (Burgess and Granato, 2007a). We did not observe any differences in SLC response frequency in either *chd7* heterozygotes or mutants (Fig. 2A,B), indicating that auditory detection is largely intact. LLC responses, however, were significantly reduced in mutants (Fig. 2C,D), suggesting that there is a specific defect within the LLC circuit.

**Figure 2.**
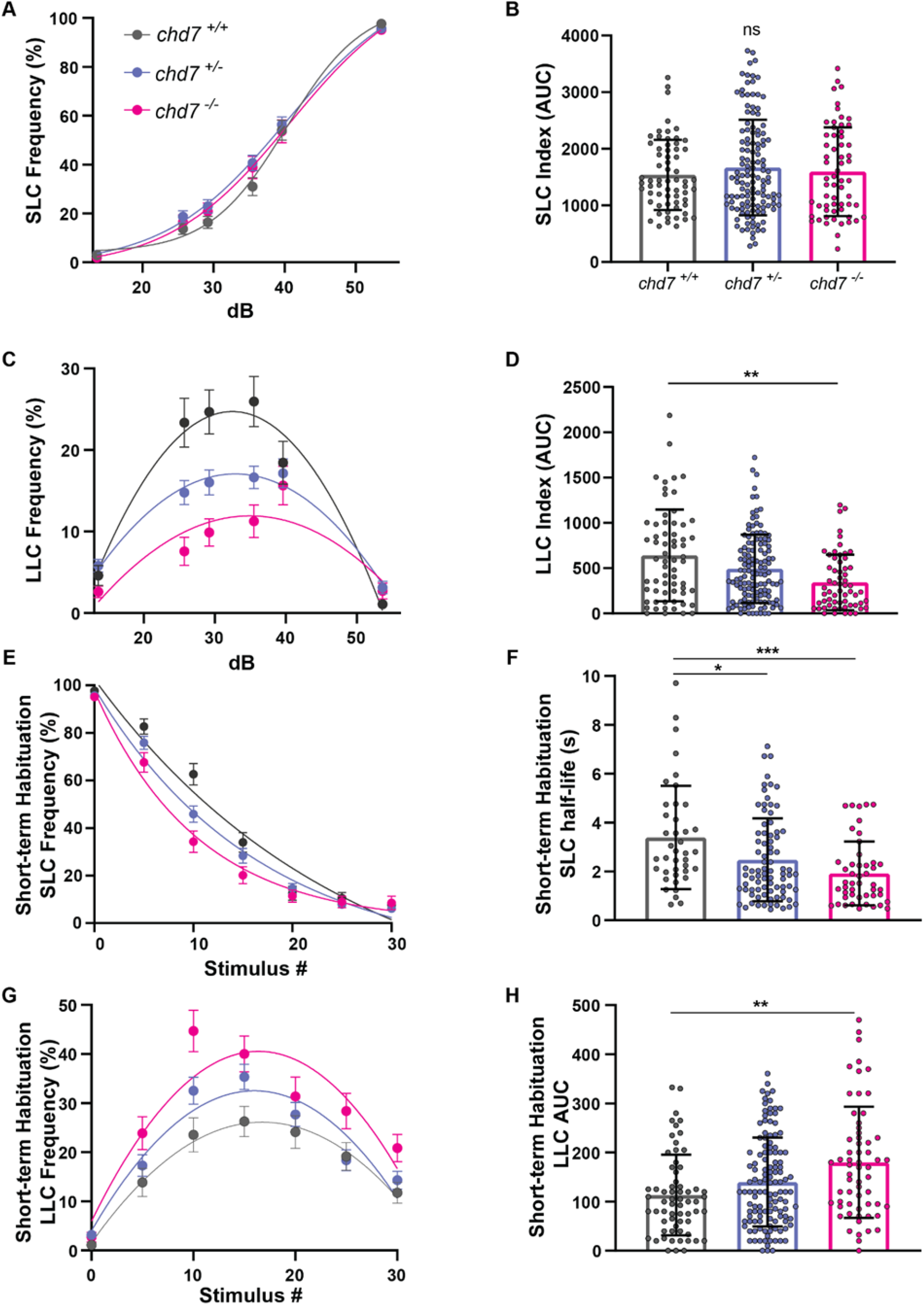
Loss of *chd7* induces a context-dependent LLC phenotype in auditory driven behaviors. **(A)** Acoustic startle responses, average short-latency c-bend (SLC) frequency as acoustic stimulus intensity increases (*chd7*^*+/+*^ n=64, *chd7* ^*+/-*^ n=127, *chd7*^*-/-*^ n=61) (mean ± SEM). **(B)** Short-latency c-bend sensitivity index, calculated by the area under the SLC frequency curves for individual larvae (mean ± SD, Ordinary one-way ANOVA with Tukey’s multiple comparison). **(C)** Long-latency c-bend (LLC) frequency as acoustic stimulus intensity increases (mean ± SEM). **(D)** Long-latency c-bend sensitivity index calculated by the area under the LLC frequency curves for individual larvae (mean ± SD, Kruskal-Wallis with Dunn’s multiple comparisons). **(E)** Short-term habituation, average SLC frequency during 30 acoustic stimuli at highest intensity (*chd7*^*+/+*^ n=40, *chd7* ^*+/-*^ n=82, *chd7*^*-/-*^ n=47) (mean ± SEM). **(F)** SLC half-life calculated by nonlinear regression (one-phase exponential decay) of SLC frequency curves for individual larvae (mean ± SD, Kruskal-Wallis with Dunn’s multiple comparisons). **(G)** Average LLC frequency during 30 acoustic stimuli at highest intensity (mean ± SEM). **(H)** LLC sensitivity index calculated by the area under the LLC frequency curves for individual larvae (mean ± SD, Kruskal-Wallis with Dunn’s multiple comparisons, *p<0.05, **p<0.01, ***p<0.001,****p<0.0001).

Following repeated strong acoustic stimulation, larval zebrafish display a simple form of non-associative learning, short-term habituation, in which SLC response frequency rapidly decreases (Roberts et al., 2011; Wolman et al., 2011). As SLCs decrease, larvae briefly shift their response bias to perform more LLCs, and eventually ignore the stimulus altogether (Jain et al., 2018). We tested whether *chd7* is required for this behavioral plasticity by presenting larvae with 30 acoustic stimuli at 53.6 dB with a 1s ISI. When we analyzed the half-life of SLC frequency curves during habituation, we found that mutants and heterozygotes habituate significantly faster than their wild-type siblings (Fig. 2E,F). Like their siblings, mutants shift their response bias toward LLCs, but this shift is exaggerated as they perform significantly more LLCs (Fig. 2G,H). These data indicate that the LLC response deficit we observed earlier following weaker, non-habituating stimuli (Fig. 2C,D) is context-dependent. Furthermore, these results suggest that *chd7* function is required to properly modulate activity in both the SLC and LLC circuits.

Having identified a defect in auditory behavior in *chd7* mutants, we next sought to determine whether this could be explained by the otolith malformations we observed (Fig. 1E, Table 1). Thus, we assessed auditory responsiveness in an independent cohort of larvae using the same 60-stimulus assay, after which all larvae were then examined for otolith defects and genotyped. As before, SLC responsiveness was normal in all groups (Fig. 3A,B), and we found that LLC responses were decreased in mutants and heterozygotes regardless of otolith defects (Fig. 3C,D). Thus, the LLC deficit is independent of ear morphology and likely arises from a defect in the central processing of auditory stimuli. To determine if motor function downstream of the SLC and LLC initiation circuits is disrupted by *chd7* loss of function, we analyzed the kinematic performance of both response types. For SLCs, latency, turn angle, and duration were normal in mutants, while the distance traveled during SLC responses was slightly reduced in mutants. For LLCs, all kinematic parameters were normal (Fig. S2E-H). These data suggest that motor function is largely normal in *chd7* mutants.

**Figure 3.**
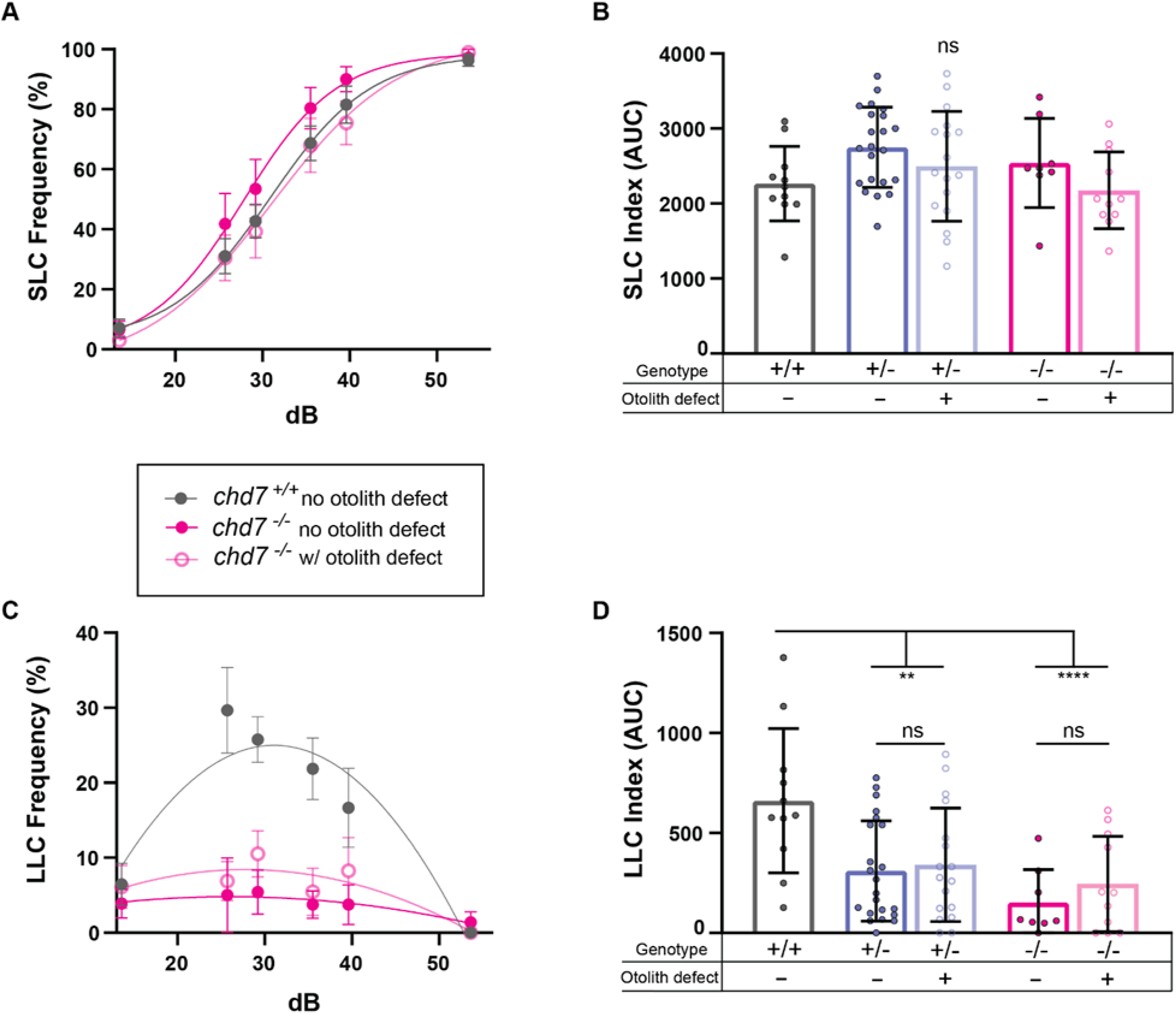
LLC deficit is independent of otolith morphology in *chd7* mutants. **(A)** Acoustic startle responses, average short-latency c-bend (SLC) frequency comparing *chd7*^*ncu101*/+^ and *chd7*^*ncu101/ncu101*^ with or without otolith defects (*chd7*^+/+^ n=11, *chd7*^+/-^ w/o otolith defect n=22, *chd7*^+/-^ with otolith defect n=17, *chd7*^-/-^ w/o otolith defect n=8, *chd7*^-/-^ with otolith defect n=11) (mean ± SEM). **(B)** Short-latency c-bend sensitivity index, calculated by the area under the SLC frequency curves for individual larvae (mean ± SD, Ordinary one-way ANOVA with Tukey’s multiple comparison). **(C)** Long-latency c-bend (LLC) frequency as acoustic stimulus intensity increases (mean ± SEM). **(D)** Long-latency c-bend sensitivity index calculated by the area under the LLC frequency curves for individual larvae (mean ± SD, Ordinary one-way ANOVA with Tukey’s multiple comparison, **p<0.01, ****p<0.0001).

We next tested the Exon 16 mutant line using the same acoustic startle assay to confirm the role of *chd7* in these auditory behaviors. As with the Exon 9 mutants, SLC response frequency was normal in heterozygotes and mutants (Fig. S3A,B). LLC response frequency, however, was not statistically significantly affected in Exon 16 mutants, although there was a downward trend (Fig. S3C,D). We also observed similar trends for short-term habituation, during which mutants’ SLCs habituated faster (Fig. S3E,F), while LLCs were increased (Fig. S3G,H). Exon 16 mutants also displayed multiple SLC kinematic defects, including increased latency, decreased turn angle, and reduced distance traveled (Fig. S4). It is possible that differences in the wild-type strain used to generate the two mutant lines may account for the phenotypic differences we observed between the two alleles. To account for this possibility, we propagated the Exon 16 allele into the TLF strain used to generate the Exon 9 allele for one generation and again tested auditory behavior in their offspring. Although less pronounced than the LLC deficit in Exon 9 mutants, we measured a significant reduction in LLC frequency in Exon 16 mutants (Fig. S5D). This data suggests that strain can influence the expression of auditory behavior phenotypes.

To further clarify the contribution of *chd7* mutations to auditory responsiveness, we performed a complementation test by crossing Exon 9 and Exon 16 heterozygotes. *chd7*^ncu101//rdu1002^ had intact SLC responses (Fig. S5E,F), and there was a downward trend in their LLC response frequency compared to wild-type siblings (Fig. S5G,H). That both Exon 9 and Exon 16 mutants show significantly reduced LLC responses when in the TLF strain while *chd7*^ncu101//rdu1002^ double heterozygotes only showed a trend indicates that there could be differential functional capacity of the two *chd7* alleles or that modifiers in the genetic background may influence the expression of the defect.

### *chd7* alleles produce varying visual and general locomotor phenotypes

Visual deficits are common in CHARGE patients (Martin et al., 2020; Onesimo et al., 2021), and therefore we assessed visual responsiveness in Exon 9 *chd7* mutants with multiple assays. First, we presented 5 dpf larvae with a series of “dark flashes” (sudden decreases in illumination) and “light flashes” (sudden increases in illumination), both of which elicit stereotyped, large-angle turns (Burgess and Granato, 2007b). Mutants performed significantly fewer dark flash responses (Fig. 4A), but their light flash responses were unaffected (Fig. 4B). This could indicate that in zebrafish, *chd7* specifically regulates the OFF retinal pathway. We also tested Exon 16 mutants for both dark flash and light flash responsiveness and found that neither behavior was significantly affected (Fig. S6A,B). This phenotypic difference between the two alleles may reflect strain differences or that the location of the induced mutation within *chd7* influences phenotype penetrance. When we tested Exon 16 mutants in the TLF background, we saw no phenotypes in either response, indicating that the visual phenotype is independent of strain (Fig. S7A,B).

**Figure 4.**
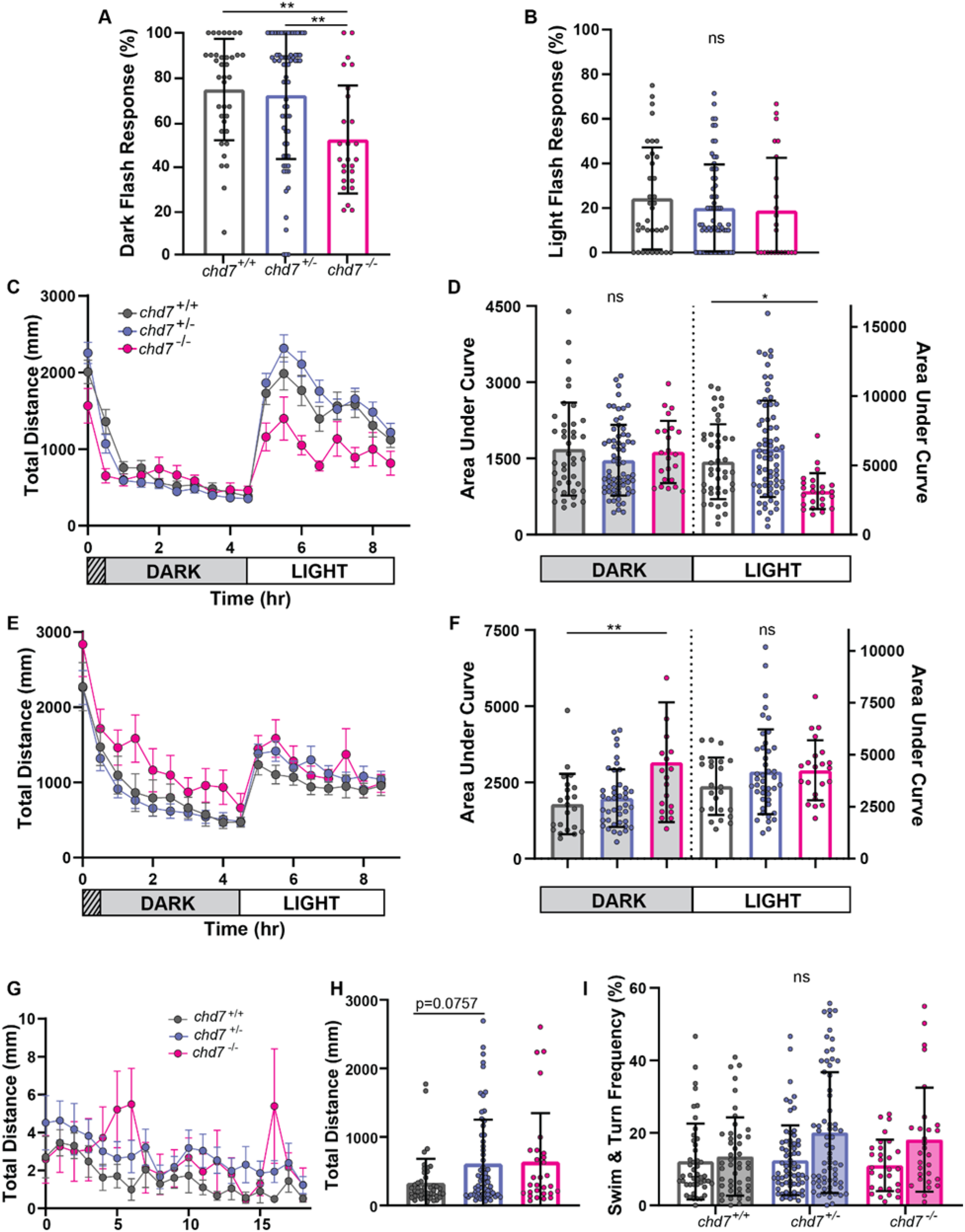
Hypo- and Hyper-activity phenotypes in Exon 9 and 16 mutants. **(A)** Exon 9 larval dark-flash and **(B)** Light-flash response frequencies (*chd7*^*+/+*^ n=37, *chd7*^+/-^ n=69, *chd7*^*-/-*^ n=26) (mean ± SD, Ordinary one-way ANOVA with Tukey’s multiple comparison, **p<0.01). **(C)** Exon 9 (*chd7*^*ncu101*^), average total distance traveled plot during 9 hours of recording, analyzed every 30 minutes, including 1 hour of acclimation in the dark, 4 hours in the dark, and 4 hours in light (*chd7*^*+/+*^ n=44, *chd7*^+/-^ n=70, *chd7*^*-/-*^ n=26) (mean ± SEM). **(D)** Exon 9, area under the curve for individual distance plots during the dark or light phase (mean ± SD, Ordinary one-way ANOVA with Dunnett’s multiple comparisons, *p<0.05,). **(E)** Exon 16 (*chd7*^*rdu1002*^), average total distance traveled (*chd7*^*+/+*^ n= 22, *chd7*^+/-^ n=42, *chd7*^*-/-*^ n=20) (mean ± SEM) and **(F)** area under the curve during the dark or light phase (mean ± SEM **(G)** Exon 9 total distance time plot during 18.5 minutes of recording (*chd7*^*+/+*^ n= 46, *chd7*^+/-^ n=67, *chd7*^*-/-*^ n=29) (mean ± SEM) **(H)** Sum of total distance traveled for individual larvae, **(I)** swim (open bars) and turn (shaded bars) frequencies (mean ± SD, Kruskal-Wallis with Dunn’s multiple comparisons).

Having identified a defect in responses to visual stimuli, we aimed to determine if the defect extends to longer-term behavior patterns. We thus tested *chd7* mutants from both the Exon 9 and Exon 16 lines in an extended visuo-motor response (VMR) assay by dark-adapting 6 dpf larvae for 30 minutes, followed by 4 hours of darkness and then 4 hours in bright light conditions (Jamadagni et al., 2021). Exon 9 mutants showed no difference in movement during the dark phase, but in the light phase showed significant hypoactivity (Fig. 4C,D). Exon 16 mutants showed a different pattern of activity, with mutants having normal activity in the light phase but hyperactivity in the dark phase (Fig. 4E,F). This result replicates the dark-phase hyperactivity recently reported in an independent *chd7* mutant line in which CRISPR/Cas9 was used to target Exon 17 (Jamadagni et al., 2021), which contributes to the same HELIc protein domain as Exon 16. We also analyzed larvae from a complementation cross of *chd7*^*ncu1019*/+^ by *chd7*^*rdu1002*/+^ adults and found no significant differences in distance traveled by double heterozygotes in both dark and light phases (Fig. S8A,B). Strain likely does not account for these differences, as larvae for these experiments, including the complementation cross, were all derived from the TLF strain. That both the Exon 9 mutants’ light-phase hypoactivity and the Exon 16 mutants’ dark-phase hyperactivity were complemented by the other allele provides further evidence that the two alleles have different functional capabilities.

Based on *chd7* mutants’ altered activity levels and kinematic defects (Fig. S2,S4), along with previous studies showing motor coordination defects, increased circling, and hyperactive behaviors in *Chd7* heterozygous mice (Alavizadeh et al., 2001; Bosman et al., 2005; Kiernan et al., 2002; Pau et al., 2004; Whittaker et al., 2017a), we recorded spontaneous movements at 50 fps for 18.5 minutes to more closely observe potential motor deficits. Exon 9 mutants showed no difference in distance moved or in the frequency of swims or turns during the assay. Exon 16 mutants, however, displayed a significant increase in swim frequency, a hyperactive phenotype consistent with their dark-phase increase in locomotion (Fig. S6E). Additionally, Exon 16 mutants derived in the TLF background also displayed an increased swim frequency (Fig. S7E), indicating a strain-independent phenotypic difference between the two alleles.

As the differences in behavioral phenotypes between the Exon 9 and Exon 16 *chd7* mutants cannot be entirely accounted for by the wild-type strain that was used, we next examined whether the differences in morphological phenotypes (Table 1, Table S1) could be explained by strain differences. Comparing Exon 9 mutants and Exon 16 mutants, we found that Exon 16 mutants did not display any otolith defects (Table S1), in stark contrast to the Exon 9 mutants (Fig. 1F, Table 1). Exon 16 mutants had a higher frequency of craniofacial defects and pericardial edema. Overall, the frequencies of mutant larvae with any type of morphological defect were similar for both alleles, but the frequency of *chd7* heterozygous larvae with morphological defects was substantially higher in Exon 9 mutants. We thus assessed morphological defects in larvae derived from a complementation cross of Exon 9^+/-^ and Exon 16^+/-^, and we observed the presence of all previously observed defects (Table S2), indicating a clear failure to complement. Together, the differences in both behavioral and morphological defects between the Exon 9 and Exon 16 mutants suggest that the location of the mutation in the *chd7* gene impacts the expression of CHARGE-related phenotypes.

### CHARGE-related phenotypes depend on *chd7* mutation location

To directly test if morphological and behavioral phenotypes in *chd7* mutants are dependent on the location of the mutation, we injected 1-cell stage TLF-strain embryos with CRISPR/Cas9 ribonucleoproteins (RNPs) using unique gRNAs targeting a site early in the gene (Exon 3), the middle of the gene (Exon 9), late in the gene (Exon 30), or a mixture of all three. The targeted regions were in the low complexity domain 12, chromodomain 2, and the low complexity domain 21, respectively. Embryos injected only with Cas9 protein served as controls. Genomic editing was detected and verified for all target sites using flanking primers for PCR followed by Sanger sequencing (Fig. S9A,B,C). At 5 dpf, larvae injected with just a single gRNA displayed lower frequencies of morphological phenotypes compared to the stable mutant lines (Table 2). Larvae injected with the 3-gRNA mixture showed similar phenotype frequencies to the stable lines, with the exception of otolith defects, which were reduced compared to Exon 9 mutants. Of the larvae injected with one gRNA, morphological phenotype frequencies were similar for Exon 9 and Exon 30 but were lowest for Exon 3. This result is contrary to a common CRISPR strategy of targeting a site earlier in the gene to produce the strongest effects.

**Table 2.** Morphological phenotype frequencies in CRISPANTS. Morphological phenotype ratio and frequencies in 5-6 dpf Cas9*/chd7* RNP*-*injected CRISPANTS.

|  | Craniofacial defect | Otolith defect | Pericardial edema | Noninflated swim bladder | Larvae with any morphological phenotype |
| --- | --- | --- | --- | --- | --- |
| Cas9 Control | - | - | - | - | 0/73, 0.0% |
| Exon 3 | - | 3/32, 9.4% | 5/32, 15.6% | 2/32, 6.3% | 11/32, 34.4% |
| Exon 9 | 2/32, 6.3% | 6/32, 18.8% | 2/32, 6.3% | 8/32, 25% | 15/32, 46.9% |
| Exon 30 | 3/40, 7.5% | 8/40, 20% | 7/40, 17.5% | 7/40, 17.5% | 15/40, 37.5% |
| Mix (3, 9, 30) | 21/42, 50% | 3/42, 7.1% | 20/42, 47.6% | 37/42, 88.1% | 39/42, 92.9% |

Finally, we tested CRISPR-injected larvae for auditory and visual behaviors as before. In contrast to the stable Exon 9 and Exon 16 lines, all four injection conditions caused a reduction in SLC responses compared to Cas9-injected controls (Fig. 5A,B). LLC responses were reduced only in larvae injected with all 3 gRNAs (Fig. 5C,D). During the short-term habituation assay all injection conditions induced faster habituation of SLC responses (Fig. 5E), and increased LLC responses during short-term habituation (Fig. 5F), similar to our findings for both the stable Exon 9 (Fig. 2) and Exon 16 lines (Fig. S1). In the long-term VMR assay, mixture-injected larvae displayed increased total distance traveled in both the dark and light phase compared to control injected siblings. Taken together, these results highlight that there are substantial phenotypic differences between CRISPR-injected larvae and larvae derived from stable mutant lines. They also further illustrate that *chd7* loss of function in zebrafish produces CHARGE-related morphological and behavioral phenotypes, providing a powerful model system to understand the biological basis for these deficits.

**Figure 5.**
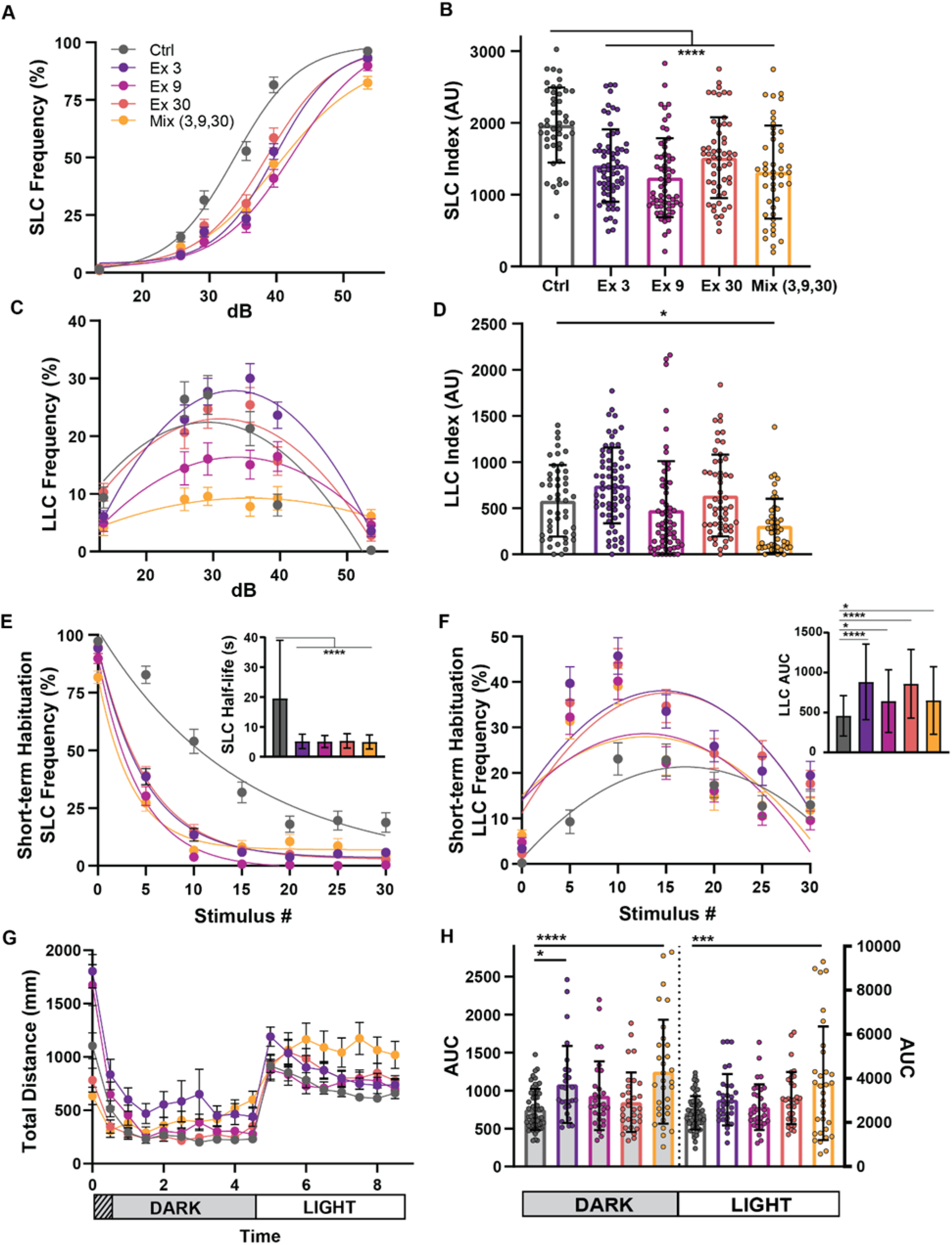
CRISPR guides targeting different *chd7* Exons induce varying phenotypes. **(A)** Acoustic startle responses, average short-latency c-bend (SLC) frequency as acoustic stimulus intensity increases (Control n=47, Exon 3 n=66, Exon 9 n=60, Exon 30 n=54, Mix n=45) (mean ± SEM). **(B)** Short-latency c-bend sensitivity index, calculated by the area under the SLC frequency curves for individual larvae (mean ± SD, Ordinary one-way ANOVA with Dunnett’s multiple comparisons). **(C)** Long-latency c-bend (LLC) frequency as acoustic stimulus intensity increases (mean ± SEM). **(D)** Long-latency c-bend sensitivity index calculated by the area under the LLC frequency curves for individual larvae (mean ± SD, Ordinary one-way ANOVA with Dunnett’s multiple comparisons). **(E)** Short-term habituation, average SLC frequency (Control n=48, Exon 3 n=61, Exon 9 n=59, Exon 30 n=57, Mix n=49) (mean ± SEM) during 30 acoustic stimuli at highest intensity with (inset) SLC half-life calculated by nonlinear regression (one-phase exponential decay) of SLC frequency curves for individual larvae (mean ± SD, Ordinary one-way ANOVA with Dunnett’s multiple comparisons). **(F)** Average LLC frequency (mean ± SEM) during 30 acoustic stimuli at highest intensity with (inset) LLC sensitivity index calculated by the area under the LLC frequency curves for individual larvae (mean ± SD, Ordinary one-way ANOVA with Dunnett’s multiple comparisons). **(G)** Average total distance traveled plot during 9 hours of recording, analyzed every 30 minutes, including 1 hour of acclimation, 4 hours in the dark, and 4 hours in light (Control n=54, Exon 3 n=27, Exon 9 n=30, Exon 30 n=31, Mix n=31) (mean ± SEM). **(H)** Area under the curve for individual larvae distance plots during the dark or light phase (mean ± SD, Ordinary one-way ANOVA with Dunnett’s multiple comparisons, *p<0.05, ***p<0.001,****p<0.000).

## Discussion

CHARGE syndrome is a heterogeneous condition with highly variable presentation that frequently includes many neurobehavioral features (Bernstein and Denno, 2005; Hartshorne and Cypher; Hartshorne et al., 2005; Hartshorne et al., 2016; Hartshorne et al., 2017; Smith et al., 2005). To better understand this heterogeneity, we generated multiple CHARGE models in zebrafish using CRISPR/Cas9 to target several locations within *CHD7*, the gene mutated in the majority of CHARGE cases (Jongmans et al., 2006; Vissers et al., 2004). Similar to the human disease, we observed variable penetrance of CHARGE-related morphological and behavioral phenotypes. Craniofacial, cardiac, visual, auditory, and locomotor defects, all of which are seen in CHARGE patients, were present in our larval zebrafish *chd7* mutants. These findings reinforce the validity of the zebrafish model for studying the pathogenesis of CHARGE syndrome, but more importantly our data provide new evidence that the location of mutations within the *CHD7* gene likely contributes to variation in the penetrance of CHARGE-related phenotypes.

Most mutations in *CHD7* leading to CHARGE syndrome are *de novo*, and they have been found throughout the gene as missense, nonsense, frameshift, splice site, and intronic mutations (Bergman et al., 2011; Jongmans et al., 2006; Legendre et al., 2017). While there is no mutational hotspot that gives rise to the disease, whether the location of mutations may contribute to the variation in phenotypes present has not been clearly defined. We thus analyzed a set of patient data with well-defined phenotypes that included their corresponding *CHD7* mutation information (Legendre et al., 2017). Because this is a relatively small patient cohort, we grouped all mutation types together based on their domain location. All intronic mutations were grouped together regardless of location. Through this analysis we found that patients with mutations in the histone-binding chromodomains presented with the highest percentage of all CHARGE phenotypes that were assessed, followed by those with mutations in the ATP helicase DEXDc domain, and the DNA-binding SANT domain (Fig. 6).

**Figure 6.**
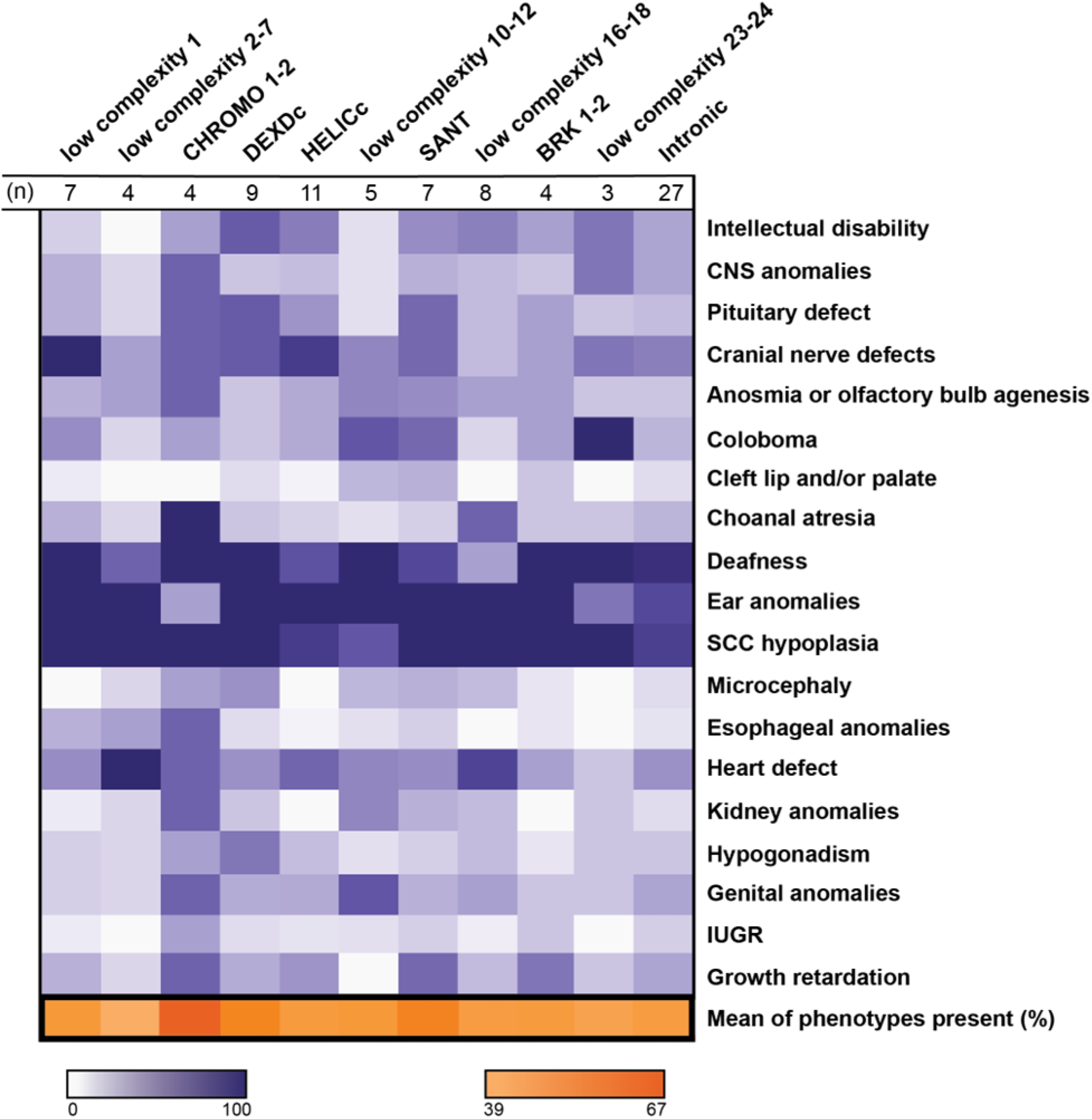
Correlation of mutation location and phenotype penetrance in a cohort of CHARGE patients. Heat map displaying the frequency of patients with a specified phenotype and a mutation within a particular *CHD7* domain; domains were determined by amino acid alignments (n=89, n= 30 patients excluded from the original pool of 119 participants due to insufficient information or mutation did not align to a domain). CHD7 domains are represented by columns and phenotypes are represented by rows. Frequencies were calculated by the number of patients with a reported phenotype and mutation in the respective domain over the total number of patients within the same domain (purples). Final row represents the overall average phenotype frequency for a specific domain (oranges). for aPatient data was obtained from the paper *Phenotype and genotype analysis of a French cohort of 119 patients with CHARGE syndrome*, Legendre et al. (2017). Darker colors indicate a higher frequency, with ranges indicated in the legends. CNS, central nervous system; SCC, semicircular canal; IUGR, intrauterine growth restriction.

This data is consistent with our findings in zebrafish larvae, in which chromodomain-targeted Exon 9 mutants (*chd7*^*ncu101*^) have higher rates of morphological phenotypes than larvae with mutations affecting the HELIc domain in Exon 16 (*chd7*^*rdu1002*^; Tables 1, S1). This includes highly penetrant otolith defects that were only observed in Exon 9 mutants. Exon 9 mutants also show more behavioral deficits than Exon 16 mutants, having reduced Mauthner cell-independent long-latency startle responses (LLCs) following acoustic stimuli, reduced O-bend responses following dark flash visual stimuli, and locomotor defects. Exon 16 mutants did show hyperactivity while in the dark, a phenotype not seen in Exon 9 mutants but one consistent with a recently described zebrafish *chd7* mutant with a different CRISPR-induced mutation in the HELIc domain (Exon 17) (Jamadagni et al., 2021). Our data also show that genetic background can contribute to variation in CHARGE-related phenotypes, as Exon 16 mutants show reduced acoustic LLC responses when maintained in the TLF strain (Fig. S5C,D), but not in the AB strain (Fig. S3C,D). Similarly, we observed a higher rate of morphological phenotypes in Exon 9 heterozygotes in the TLF strain (Table 1) than in those from a complementation cross to the AB-background HELIc domain line (Table S2). Thus, the AB background may contain genetic modifiers that partially suppress some CHARGE-related phenotypes. Hyperactivity, however, was observed in Exon 16 mutants independent of strain (Fig. 4, Fig. S6,S7), but not in Exon 9 mutants, highlighting the domain dependence of some behavioral phenotypes.

The domain-dependence of some CHARGE-related phenotypes also extended to our CRISPR-injected F_0_ larvae. Those injected with CRISPR/Cas9 targeting the chromodomain 2 (Exon 9) had greater overall penetrance of developmental phenotypes such as craniofacial, otolith, and pericardial defects than those in which low complexity domains were targeted (Exon 3: low complexity 14; Exon 30: low complexity 24; Table 2). This pattern was not seen for the behavioral phenotypes we assessed, however, as larvae injected with a single gRNA had largely similar behavioral profiles regardless of the *chd7* target site. Larvae in which all three gRNAs were injected together showed the highest penetrance of both morphological and behavioral phenotypes, with 93% of injected larvae having at least one morphological defect, along with significant defects in both Mauthner-dependent SLCs and Mauthner-independent LLCs and hyperactivity in both the dark and light phases of the visuo-motor response assay (Fig. 5). The F_0_ larvae differed in some respects from *chd7* mutants derived from our stable mutant lines, such as the SLC deficit that was not observed in stable mutants. As the same wild-type strain was used for all of the F_0_ larvae, these phenotypic differences are likely due to the mosaicism of the F_0_ larvae or perhaps differences in genomic adaptation to the induced mutations between acutely injected larvae and those inheriting a single stable mutation. It is important to note that the early-exon targeted larvae (Exon 3) showed the fewest phenotypes, as targeting early exons with CRISPR/Cas9 is a frequent strategy for attempting to create null alleles in addition to using multiple guides (Hoshijima et al., 2019). Our data demonstrate that *chd7* mutations not only produce variable phenotypes, but that the location of the targeted site, even if a premature stop codon is created, contributes to the expression of CHARGE-related phenotypes.

Our larval zebrafish CHARGE models recapitulate disease phenotypes seen previously in multiple animal systems, including *C. elegans, Drosophila, Xenopus*, zebrafish, and mouse. We summarized these CHARGE models and their primary phenotypes in Supplemental Table 3. In *C. elegans*, loss of *chd7* via RNA interference causes undetached cuticle, gonad migration defects, and stunted length (Jofré et al., 2022). Although we did not find significant differences in length and size in our zebrafish *chd7* mutants, mouse models have been reported to have delayed growth (Bosman et al., 2005), a symptom sometimes seen in CHARGE patients pre- and postnatally (Antoniou et al., 2019; Kim et al., 2022; Traisrisilp et al., 2021). In *Drosophila*, mutations in *kismet*, the orthologue of *chd7*, cause movement deficits, eye defects, and wing abnormalities, as well as improper axon pruning, mislocalized neurons, and decreased glutamate receptor localization (Ghosh et al., 2014; Melicharek et al., 2010). These defects in neural architecture in *kis*^*-/-*^ may provide insight into potential mechanisms driving the deficits we see in our zebrafish model. However, *kismet* is an orthologue for both *chd7* and *chd8* in vertebrates and thus plays additional functional roles, including facilitation of transcriptional elongation (Srinivasan et al., 2005). Additionally, the reported alleles in *kis*^*-/-*^ affect Exon 3, which contributes to most of the n-terminus of the protein including the chromodomains and ATP-helicase domain, making it difficult to compare phenotypes and potential domain dependence. *chd7* knock-down with a translation-blocking morpholino in *Xenopus* tadpoles resulted in similar morphological phenotypes to those in our zebrafish larvae, including craniofacial, heart, and otolith defects (Wysocka et al., 2010). Finally, prior larval zebrafish CHARGE models have largely used morpholinos to knock down *chd7*, and these morphants display pericardial edema, craniofacial defects, and otolith defects (Asad et al., 2016; Balow et al., 2013; Liu et al., 2018; Patten et al., 2012), similar to our Exon 9 mutants. *Chd7* morphants also have neural crest cell migration defects (Asad et al., 2016), reduced T-cell proliferation (Liu et al., 2018), and disorganized retinal organization (Patten et al., 2012). More recent CRISPR-generated *chd7* mutant larvae display similar morphological phenotypes to those in the present study including pericardial edema, craniofacial defects (Jamadagni et al., 2021), and uninflated swim bladders (Cloney et al., 2018), in addition to defects in gastrointestinal motility (Cloney et al., 2018). Together, these studies underscore the highly conserved and multiple roles of *chd7* in development.

In mice, homozygous *CHD7* mutations are embryonically lethal, so research has been largely limited to *CHD7*^*+/-*^ derived from ENU mutagenesis (Bosman et al., 2005; Hrabé de Angelis et al., 2000; Kiernan et al., 2002; Nolan et al., 2000). *CHD7*^*+/-*^ mice, like our zebrafish mutants, display ear and cardiovascular abnormalities (Table S3) (Bosman et al., 2005). *CHD7*^*+/-*^ also exhibit circling behaviors (Bosman et al., 2005), an indication of vestibular dysfunction, and reduced olfactory function (Bergman et al., 2010). Cerebellar-targeted *CHD7* conditional mutants show motor delays and coordination deficits (Whittaker et al., 2017a). We did not see evidence of vestibular or coordination deficits in our zebrafish models, and so these phenotypes could indicate unique roles for mammalian *CHD7*, but further analysis is needed as we did not examine limb movement in our locomotor assays due to the difficulty of imaging the movement of the pectoral fins in 5 dpf larvae. As with our zebrafish mutants and CHARGE patients, the CHARGE-related phenotypes in these mouse models are highly variable, including variation in the penetrance and severity of ear phenotypes between the multiple *CHD7* alleles with mutations in different parts of the gene (Bosman et al., 2005; Kiernan et al., 2002). Within one of these lines, *Whirligig*, phenotypes also had variable penetrance, with abnormalities in the ear and female external genitalia being highly penetrant (>90%), while eye, cardiovascular, palate, and olfactory defects having reduced penetrance (<50%) (Bergman et al., 2010; Bergman et al., 2011), despite using an inbred mouse strain. While zebrafish strains are fairly outbred and maintain significant genetic diversity (Balik-Meisner et al., 2018; Butler et al., 2015), our data also show variable penetrance of phenotypes within a single line. Furthermore, multiple sibling pairs with CHARGE syndrome that carry the same deleterious mutation in *CHD7* have been identified, with phenotypic differences between siblings (Jongmans et al., 2006). Therefore, the *CHD7* mutation location and genetic background are not the only factors that give rise to phenotypic differences, and variation in the fetal microenvironment and developmental stochasticity also likely contribute to the heterogeneity of CHARGE phenotypes (Bergman et al., 2011).

The diverse roles of *CHD7* in the development of multiple tissues make for a challenge in determining the precise cellular and molecular pathways leading to each of the many CHARGE phenotypes. In the nervous system alone, *CHD7* promotes neuronal differentiation (Feng et al., 2013; Feng et al., 2017; He et al., 2016; Jones et al., 2015), is required for proper neurite development (Yao et al., 2020), regulates cerebellar organization through *RELN* (Whittaker et al., 2017a), promotes GABAergic neuron development through *paqr3* (Jamadagni et al., 2021) and is required for neural crest cell differentiation and migration (Fan et al., 2021; Okuno et al., 2017), thereby contributing to multiple disease manifestations such as cranial nerve, cardiac, and craniofacial defects (Wysocka et al., 2010). Our Exon 9 mutants display both morphological defects in the ear and a deficit in auditory LLC responses (Fig. 1, 2). We found that the LLC deficit is independent of ear morphology, indicating that *chd7* likely regulates this behavior centrally (Fig. 3). Defects in the auditory nerve could cause this phenotype, although the fact that SLC responses are normal following non-habituation acoustic stimuli suggests that the defect may be more specific to the LLC circuit, which includes a set of pre-pontine neurons in rhombomere 1 (Marquart et al., 2019). During short-term habituation, however, SLC responses decrease faster and LLC responses are increased in *chd7* mutants (Fig. 2E-H), indicating that *chd7* also regulates the function of neurons that modulate these acoustic startle circuits. It is possible that abnormal GABAergic neuron development could underlie these defects (Jamadagni et al., 2021), as inhibitory input is crucial for normal habituation (Marsden and Granato, 2015).

In conclusion, our results provide additional support for the highly conserved roles of *CHD7* in the development of many tissues and processes, including the nervous system and behavior. Our findings underscore the highly heterogeneous nature of CHARGE phenotypes across model systems, but we also show new evidence that the location of mutations within *CHD7* contributes to variation in disease-related phenotypes. Finally, our new CHARGE models create opportunities for future efforts to take advantage of the larval zebrafish system to define brain-wide roles for *CHD7*, identify genetic modifiers of CHARGE-related phenotypes, and screen for compounds that may restore function in pre-clinical models of CHARGE.

## Materials and Methods

### *Danio rerio* Husbandry and Maintenance

All animal use and procedures approved by the North Carolina State University Institutional Animal Care and Use Committee. *chd7* CRISPANTS and Exon 9 line (*chd7* ^ncu101/+^) were maintained in the *Tüpfel long fin* (TLF) background. The TLF strain originated from University of Pennsylvania stocks. The Exon 16 line (*chd7* ^rdu1002/+^), gifted by Dr. Erica Davis from Northwestern University, was maintained in the AB and TLF background. Adult zebrafish were housed in 5 fish/L density under a 14 hr:10hr light:dark cycle at ∼28°C, and were fed rotifers, *artemia* brine shrimp, and GEMMA (Skretting).

To generate embryos for larval testing, male and female pairs were placed in mating boxes (Aquaneering) containing system water and artificial grass. 1-2 hours into the subsequent light cycle of the following day, embryos were collected and placed into petri dishes containing 1x E3 embryo media. Embryos were sorted for fertilization under a dissecting scope at ∼6 hours post fertilization (hpf) and placed into petri dishes with n ≤65. All embryos were reared in a temperature-controlled incubator at 29°C on a 14h:10h light dark cycle. Each day until testing, a 50-75% media change was performed, and any dead larvae were removed. During survival analysis, dead larvae were saved for future genotyping analysis. All experiments were performed blind to genotype.

### Generation of *chd7* ^*ncu101*/+^ and *chd7* CRISPANTS

Single guide RNAs (Supplemental Table. 4) were designed using CHOPCHOP (https://chopchop.cbu.uib.no/) and CRISPRscan (https://www.crisprscan.org/) using the zebrafish reference genome (GRCz11). Microinjections in one-cell stage embryos were performed with an injection mix containing a 50 µM gRNA duplex, Cas9 protein, 0.5% phenol red, and nuclease-free H2O. Embryos were generated from crossing TLF wild-type pairs. *Chd7* mutagenesis was confirmed in injected larvae using the IDT genomic cleavage assay detection kit with T7 endonuclease digestion, and Sanger sequencing.

### DNA extraction and Genotyping via PCR

Larvae were individually placed in 96-well plates containing methanol (MeOH) for tissue fixation. DNA was extracted from whole larvae using a lysis buffer of 25mM sodium hydroxide, 0.2 mM EDTA (base solution), and 40mM Tris-HCl (neutralization solution). For the Exon 9, 7 bp deletion mutation forward and reverse primers amplified a 289 bp fragment (wild-type). Amplicons were resolved on a 3% agarose gel using gel electrophoresis. For the Exon 16, 1bp deletion mutation forward and reverse primers amplified a 349 bp fragment (wild-type). The 1bp deletion created a unique restriction site, and amplicons were digested with *XcmI* and resolved on a 1% agarose gel (Table S5).

### Morphology and Survival Assessments

#### Morphological Phenotypes and Brightfield imaging

Larvae were assessed for morphological phenotypes at 5-6 dpf and imaged using brightfield microscopy (Nikon SMZ25). Larvae were laterally mounted in ∼1% low-melting point agarose in 1X E3 medium. Images were acquired at 10X total magnification, and phenotypes were scored following image capture. Larvae were unmounted and individually placed in 96-well plates containing MeOH for tissue fixation, DNA extraction, and genotyping. Images were masked from genotype during assessment, and manually scored by multiple experimenters.

#### Survivability

For survival assessments, larvae were tracked daily from fertilization sort (∼6 hpf) until 30 days post fertilization. Larvae were reared in system water by 7 dpf and fed according to the lab’s SOP. Fish were checked at the same time of day, and dead fish were removed and individually placed in MeOH for future genotyping. At 30 days, the remaining juveniles were euthanized and processed for genotyping.

#### Behavioral Assays and Analyses

Larvae were tested at 5-6 days post fertilization (dpf) within their normal light cycle. Larvae were screened for severe morphological defects affecting mobility including large pericardial edema and bent/missing tails. Swim bladder inflation and less severe defects (those not affecting mobility) were ignored. Post-screening, larvae adapted to ambient testing-rig lighting for 30 minutes before placement in a 6×6 well grid containing 36 test arenas (9 mm in diameter). Larvae were covered in equal amounts of 1X E3 medium in each well (∼200 uL). The testing arena was a custom laser-cut 2 mm (depth) clear acrylic square adhered to a 3 mm white acrylic square and anchored to an aluminum rod (6 × 1 × 1 cm) for attachment to the acoustic shaker (Bruel and Kjaer). Behavioral assays including spontaneous movement, acoustic startle, short-term habituation, dark-flash, and light-flash were tracked and analyzed using FLOTE-software as previously described (Burgess and Granato, 2007a; Burgess and Granato, 2007b; Marsden et al., 2018). The long-term locomotor behavioral assay was performed using the DanioVision (Noldus) apparatus and analyzed using EthoVision XT video tracking (Noldus).

#### Spontaneous Movement

Once placed into arenas, larvae acclimated for 3 minutes and subsequently recorded for 18.5 minutes at 50 frames per second (fps), 640×640 resolution.

### Acoustic Startle Response and Auditory behaviors

Larvae acclimated for 2 minutes prior to auditory stimulus onset. Larvae received a total of 60 acoustic stimuli, 10 randomized trials of 6 intensities (13.6 dB, 25.7, 29.2, 35.5, 39.6, 53.6), with a 20s inter-stimulus interval (ISI). Immediately following the acoustic startle response assay, larvae received 30 acoustic stimuli (53.6 dB) with a 1s ISI to induce short-term habituation. Recordings were captured at 120 frames per stimulus, 1000 fps, and 640×640 resolution. Following tracking and analysis, individual larval responses were filtered by response frequency (% React) during the loudest acoustic stimuli (≥ 70%), and the number of times larvae were tracked across each trial of 10 (≥7) for the acoustic startle response and each trial of 5 (≥3) for short-term habituation.

### Visual Startle Responses, Dark-Flash/ Light-Flash

Larvae acclimated in white light via an LED light (0.4 V) for 3 minutes prior to dark-flash onset. Larvae received a total of 10 “dark-flashes” or 1s intervals of the light turning off with a 30s ISI. Next, larvae received a total of 10 “light-flashes” or 1s intervals of brighter light (10V). Recordings were captured at 1000 frames per stimulus, 1000 fps, and 640×640 resolution.

### RNA extraction and cDNA synthesis

For qPCR analysis, 5 dpf larvae were anesthetized using 0.2% Tricaine in 1X E3 medium. Larval heads were dissected using a 15°, 5mm scalpel, briefly washed in 1X PBS, and placed in a 96-well plate containing ∼200uL *RNALater* (Invitrogen). Remaining tissue was placed in a corresponding well containing MeOH for DNA extraction and genotyping. Heads were briefly chilled on ice and placed in 4°C overnight for *RNAlater* permeation. Following ∼8 hours incubation, heads were moved to -20°C. Once genotypes were obtained, wild-type, heterozygous, and mutant heads were pooled for RNA extraction (3 biological replicates per genotype). Tissue was homogenized using Trizol, and RNA was extracted using RNeasy Mini kits (Qiagen, ID: 217084) according to manufacture protocol. For cDNA synthesis, 1 ng-5 μg of RNA per reaction was used in SuperScript™ First-Strand synthesis kits (Invitrogen, ID: 11904018). Nucleic acid purity was assessed using a NanoPhotometer® (Implen).

### qPCR

qPCR was performed with SYBR® Green PCR master mix. For *chd7*, forward and reverse intron-spanning primers annealed downstream of the mutation targets in Exon 9 (*chd7*^*ncu101/*^) and Exon 16 (*chd7*^*rdu1002*^*/)* alleles. *Eif1a* was the housekeeping gene used for quality control and normalizing *chd7* expression data.

### Statistics

Statistical analyses were completed using Prism 8 (GraphPad). All datasets were tested for normality using the Shapiro-Wilk test. ANOVA and multiple comparisons, or Kruskal Wallis, nonparametric tests and Dunn’s multiple comparisons were used, respectively. For all experiments and data sets, the p-value threshold was set to α=0.05. All data are represented as mean ± standard deviation, unless otherwise noted in the figure legends.

## Acknowledgments

We would like to thank Dr. Erica Davis (Northwestern University, Feinberg School of Medicine) for supplying the *chd7*^*rdu1002*/+^ line. We are also grateful to Rachael Bieler for additional proofreading help with the manuscript.

## Data availability

All data is available upon request.

## Author contributions

Project conceptualization: K.C.M., D.R.H.; Methodology: D.R.H., K.C.M.; Investigation: D.R.H., P.M.L., A.A.B., D.F.B., K.C.M.; Data curation: D.R.H., P.M.L., A.A.B.; Writing: D.R.H., P.M.L., K.C.M; Writing-review & editing: D.R.H., K.C.M.; Supervision: K.C.M.; Funding acquisition: K.C.M.

