## Supplemental information for "Morphological and sensorimotor phenotypes in a zebrafish CHARGE syndrome model are domain-dependent"

A

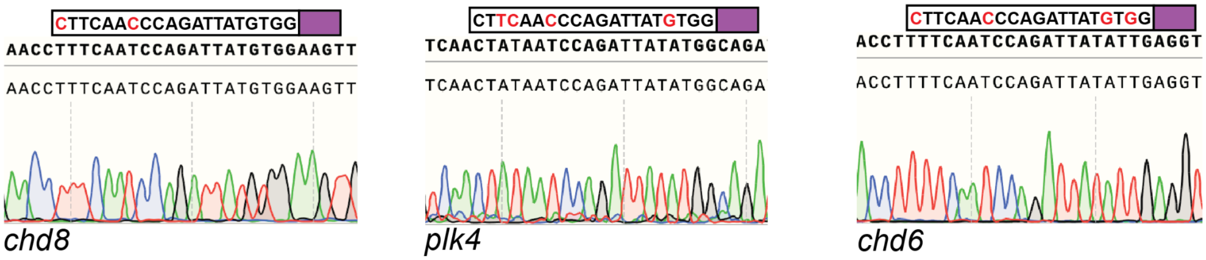

**Supplemental Figure 1. *chd7* guide RNA does not induce off-target effects in Exon 9 line.**

**(A)** Example chromatograms of top three predicted off-target sites from Exon 9 (*chd7*$^{ncu102/+}$) gRNA in *chd8, plk4,* and *chd6.* Guide RNA sequence included with nucleic acid mismatches in red, and PAM site highlighted with purple box.

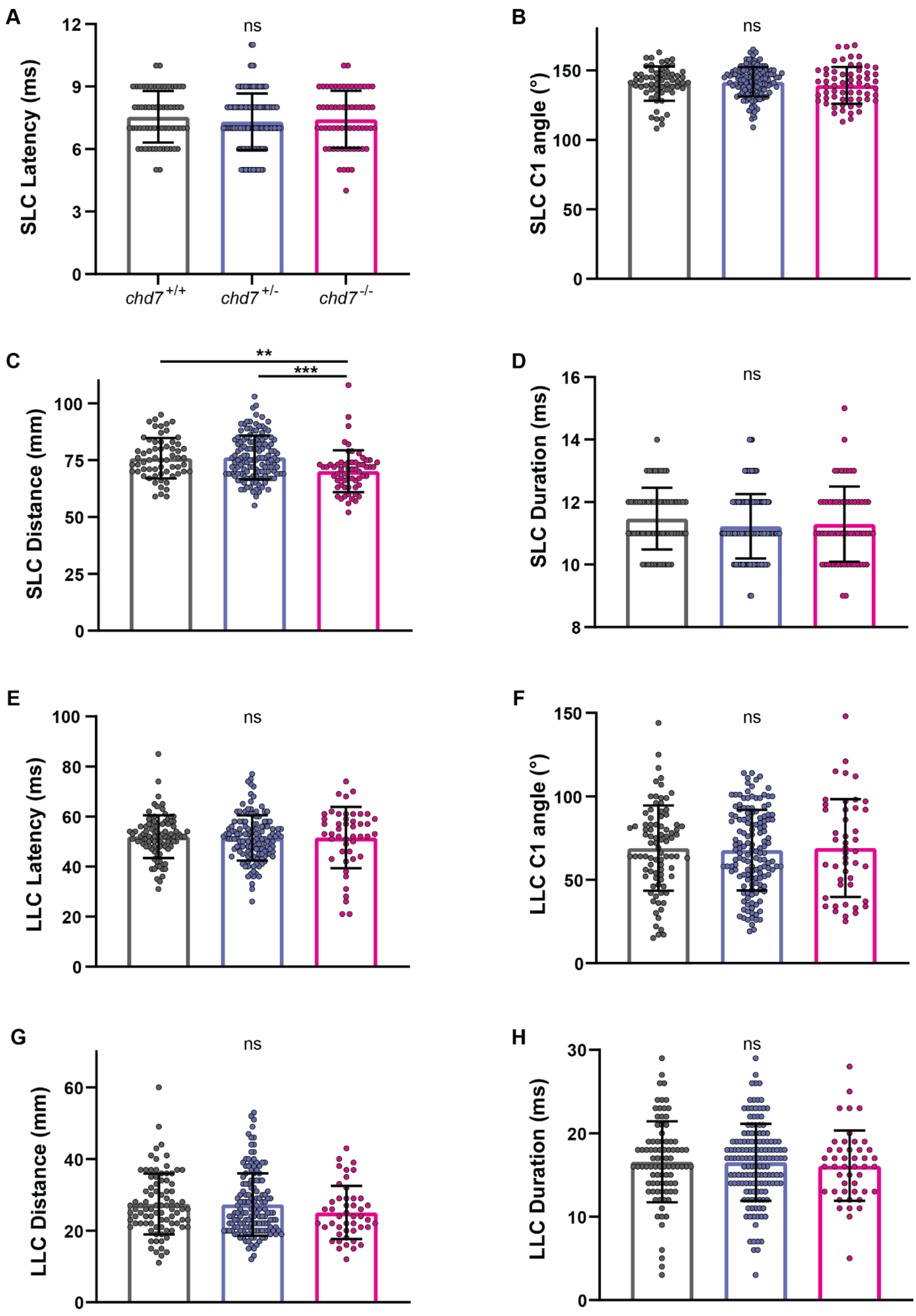
A
ns
SLC Latency (ms)
chd7 +/+
chd7 +/-
chd7 -/-
B
ns
SLC C1 angle (°)
C
**
***
SLC Distance (mm)
D
ns
SLC Duration (ms)
E
ns
LLC Latency (ms)
F
ns
LLC C1 angle (°)
G
ns
LLC Distance (mm)
H
ns
LLC Duration (ms)

**Supplemental Figure 2. Acoustic startle kinematics is intact in Exon 9 mutants. (A)** SLC response kinematics including latency of response initiation, **(B)** C1 bend angle, **(C)** average distance traveled, and **(D)** duration of SLC response (mean ± SD, Kruskal-Wallis with Dunn's multiple comparisons, $^{**}p<0.01$, $^{***}p<0.001$) **(E)** LLC response kinematics including latency, **(F)** bend angle, **(G)** average distance (mean ± SD, Kruskal-Wallis with Dunn's multiple comparisons), and **(H)** duration (mean ± SD, Ordinary one-way ANOVA with Tukey's multiple comparison).

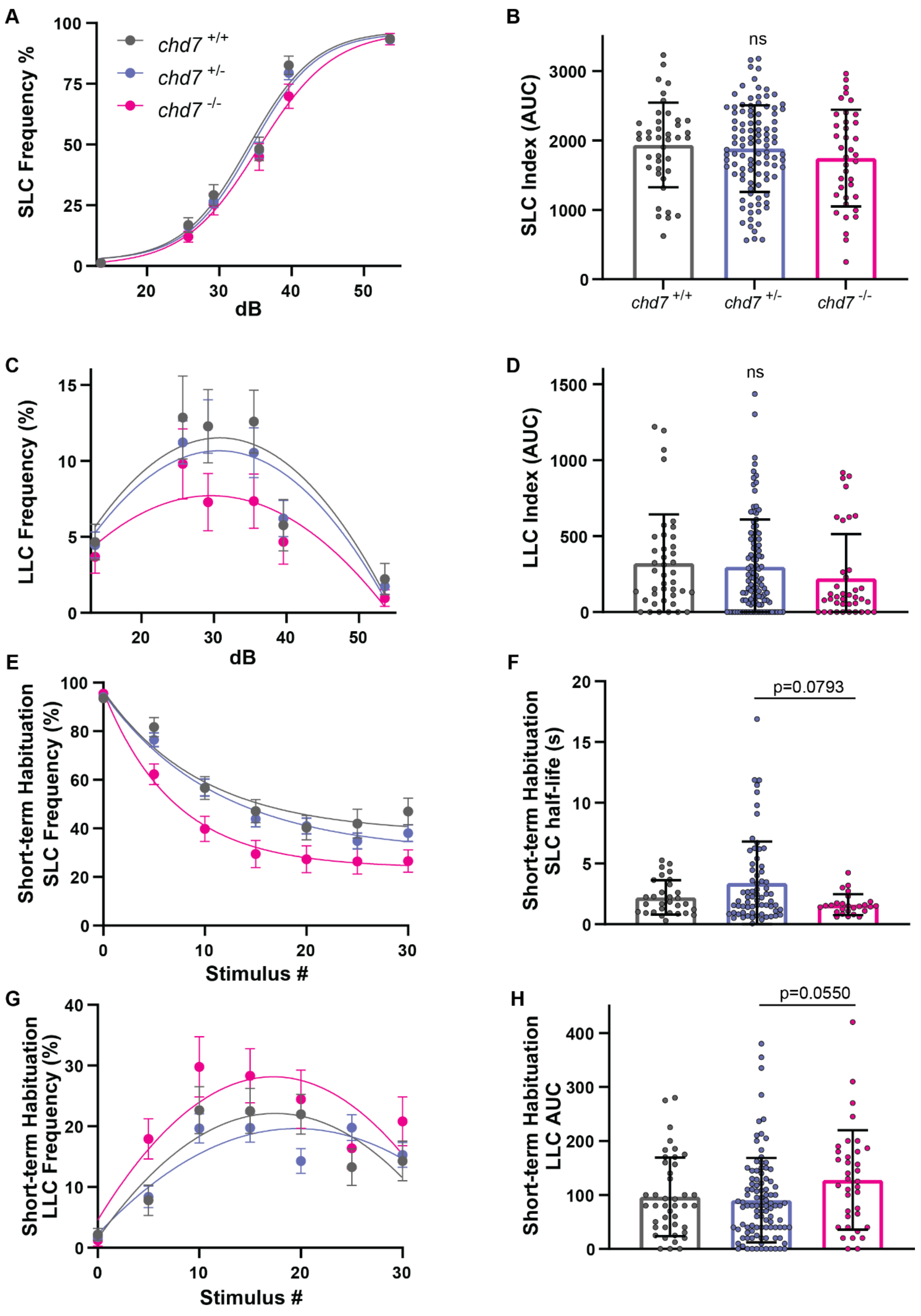
A
SLC Frequency %
dB
chd7 +/+
chd7 +/-
chd7 -/-
B
SLC Index (AUC)
ns
chd7 +/+
chd7 +/-
chd7 -/-
C
LLC Frequency (%)
dB
D
LLC Index (AUC)
ns
E
Short-term Habituation SLC Frequency (%)
Stimulus #
F
Short-term Habituation SLC half-life (s)
p=0.0793
G
Short-term Habituation LLC Frequency (%)
Stimulus #
H
Short-term Habituation LLC AUC
p=0.0550

**Supplemental Figure 3. Exon 16 mutants have intact acoustic startle responses.**

**(A)** Acoustic startle responses, average short-latency c-bend (SLC) frequency as acoustic stimulus intensity increases (*chd7*$^{+/+}$ n=60, *chd7* $^{+/-}$ n=131, *chd7*$^{-/-}$ n=54) (mean ± SEM). **(B)** Short-latency c-bend sensitivity index, calculated by the area under the SLC frequency curves for individual larvae (mean ± SD, Ordinary one-way ANOVA with Tukey's multiple comparison). **(C)** Long-latency c-bend (LLC) frequency as acoustic stimulus intensity increases (mean ± SEM). **(D)** Long-latency c-bend sensitivity index calculated by the area under the LLC frequency curves for individual larvae (mean ± SD, Kruskal-Wallis with Dunn's multiple comparisons). **(E)** Short-term habituation, average SLC frequency during 30 acoustic stimuli at highest intensity (*chd7*$^{+/+}$ n=30, *chd7* $^{+/-}$ n=67, *chd7*$^{-/-}$ n=26) (mean ± SEM). **(F)** SLC half-life calculated by nonlinear regression (one-phase exponential decay) of SLC frequency curves for individual larvae (mean ± SD, Kruskal-Wallis with Dunn's multiple comparisons). **(G)** Average LLC frequency during 30 acoustic stimuli at highest intensity (*chd7*$^{+/+}$ n=42, *chd7* $^{+/-}$ n=101, *chd7*$^{-/-}$ n=36) (mean ± SEM). **(H)** LLC sensitivity index calculated by the area under the LLC frequency curves for individual larvae (mean ± SD, Kruskal-Wallis with Dunn's multiple comparisons).

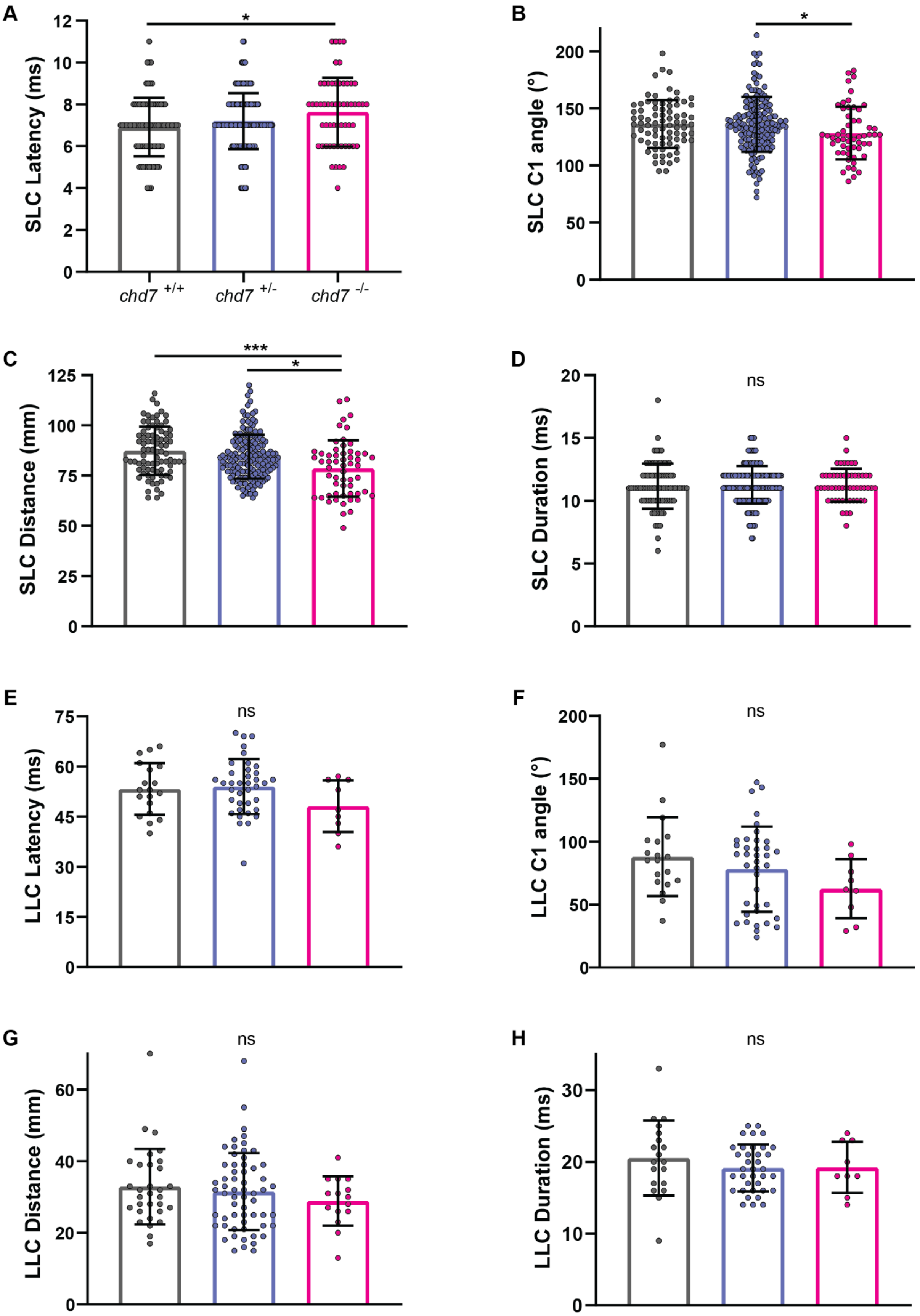
A
SLC Latency (ms)
chd7 +/+
chd7 +/-
chd7 -/-
B
SLC C1 angle (°)
C
SLC Distance (mm)
D
SLC Duration (ms)
ns
E
LLC Latency (ms)
ns
F
LLC C1 angle (°)
ns
G
LLC Distance (mm)
ns
H
LLC Duration (ms)
ns

**Supplemental Figure 4. Acoustic startle kinematics in exon 16 mutants. (A)** SLC response kinematics including latency of response initiation, **(B)** size of C1 bend angle, **(C)** average distance traveled, and **(D)** duration of SLC response (mean ± SD, Kruskal-Wallis with Dunn's multiple comparisons, *p<0.05, ***p<0.001) **(E)** LLC response kinematics including latency (mean ± SD, Ordinary one-way ANOVA with Tukey's multiple comparison), **(F)** bend angle, **(G)** average distance, and **(H)** duration (mean ± SD, Kruskal-Wallis with Dunn's multiple comparisons).

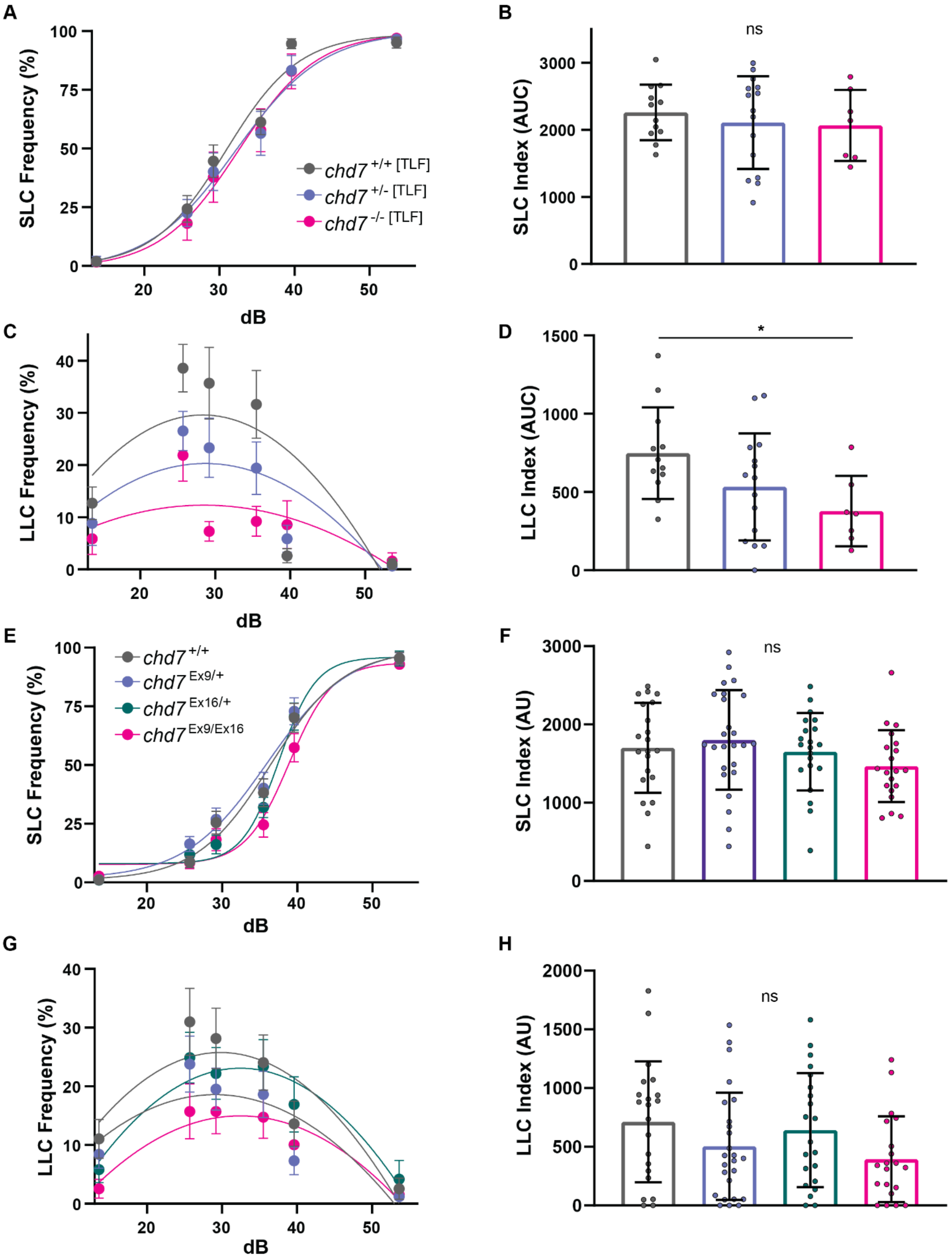
A
SLC Frequency (%)
dB
chd7 +/+ [TLF]
chd7 +/- [TLF]
chd7 -/- [TLF]
B
SLC Index (AUC)
ns
C
LLC Frequency (%)
dB
D
LLC Index (AUC)
*
E
SLC Frequency (%)
dB
chd7 +/+
chd7 Ex9/+
chd7 Ex16/+
chd7 Ex9/Ex16
F
SLC Index (AU)
ns
G
LLC Frequency (%)
dB
H
LLC Index (AU)
ns

**Supplemental Figure 5. Zebrafish strain and mutation type affect the LLC phenotype in *chd7* mutants. (A)** Acoustic startle responses of Exon 16 mutants propagated in the TLF background, average short-latency c-bend (SLC) frequency as acoustic stimulus intensity increases (*chd7*$^{+/+}$ n=12, *chd7* $^{+/-}$ n=15, *chd7*$^{-/-}$ n=7) (mean ± SEM). **(B)** Short-latency c-bend sensitivity index, calculated by the area under the SLC frequency curves for individual larvae (mean ± SD, Ordinary one-way ANOVA with Tukey's multiple comparison). **(C)** Long-latency c-bend (LLC) frequency as acoustic stimulus intensity increases (mean ± SEM). **(D)** Long-latency c-bend sensitivity index calculated by the area under the LLC frequency curves for individual larvae (mean ± SD, Ordinary one-way ANOVA with Tukey's multiple comparison, *p<0.05). **(E)** Acoustic startle responses of double heterozygotes propagated from a complementation cross, average short-latency c-bend (SLC) frequency (*chd7*$^{+/+}$ n=21, *chd7*$^{ncu101/+}$ n=25, *chd7*$^{rdu1002/+}$ n=21, *chd7*$^{ncu101/rdu1002}$ n=20) (mean ± SEM). **(F)** SLC sensitivity index (mean ± SD, Ordinary one-way ANOVA with Tukey's multiple comparison). **(G)** Long-latency c-bend (LLC) frequency (mean ± SEM) and **(H)** LLC sensitivity index (mean ± SD, Kruskal-Wallis with Dunn's multiple comparisons).

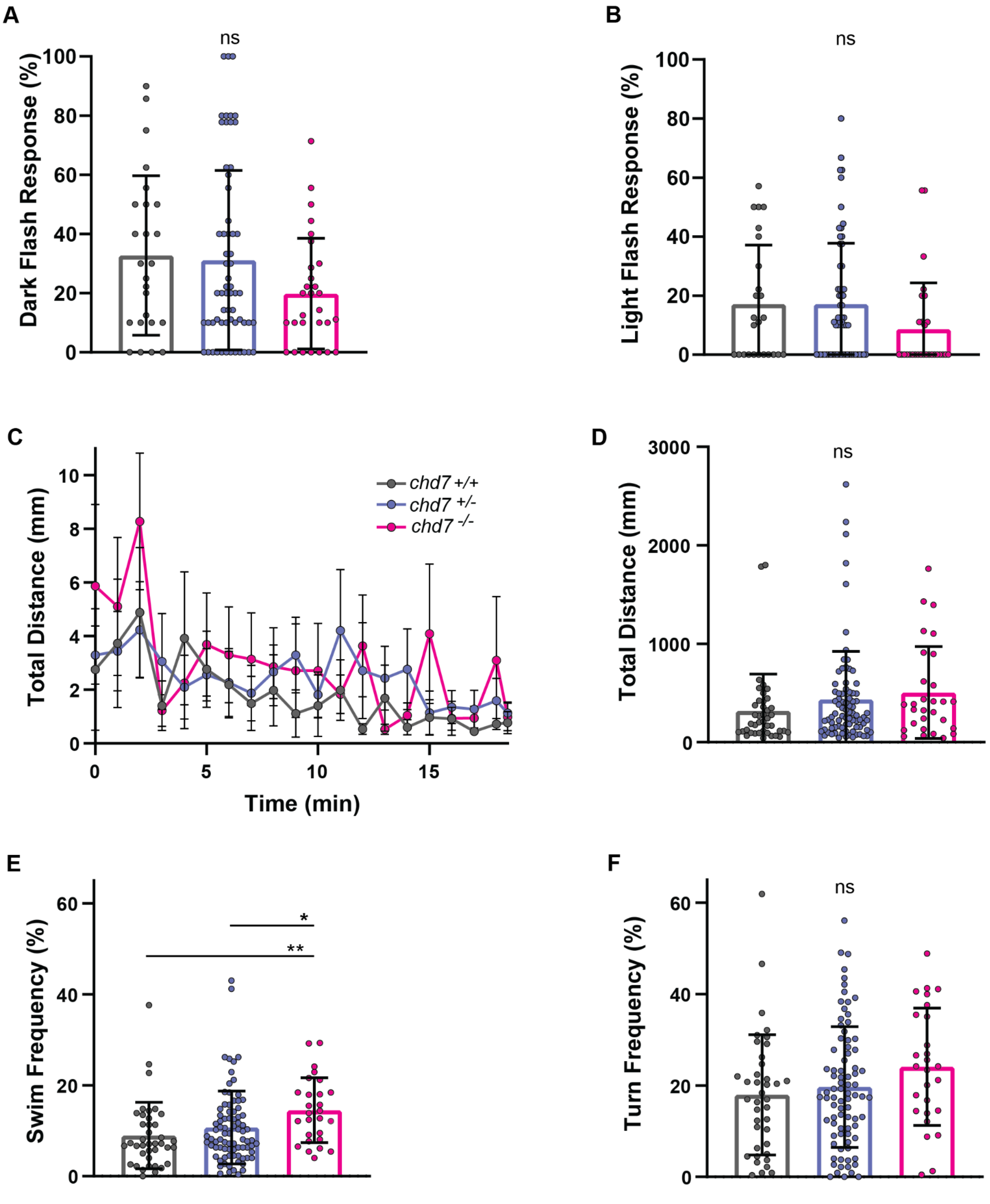

A
ns
Dark Flash Response (%)
100
80
60
40
20
0
B
ns
Light Flash Response (%)
100
80
60
40
20
0
C
Total Distance (mm)
10
8
6
4
2
0
0
5
10
15
Time (min)
chd7 +/+
chd7 +/-
chd7 -/-
D
ns
Total Distance (mm)
3000
2000
1000
0
E
*
**
Swim Frequency (%)
60
40
20
0
F
ns
Turn Frequency (%)
60
40
20
0

**Supplemental Figure 6. General locomotor and visual behaviors in Exon 16 mutants. (A)** Dark flash and **(B)** Light flash response frequencies (mean ± SD, Kruskal-Wallis with Dunn's multiple comparisons) **(C)** Average total distance traveled plot during 18.5 minutes of recording, represents one testing clutch for *chd7*$^{+/+}$*, chd7*$^{rdu1002/-}$*,* and *chd7*$^{rdu1002/rdu1002}$ (*chd7*$^{+/+}$ n= 39, *chd7*$^{+/-}$ n=82, *chd7*$^{-/-}$ n=27) (mean ± SEM). **(D)** Sum of total distance traveled for individual larvae, **(E)** swim and **(F)** turning frequencies (mean ± SD, Kruskal-Wallis with Dunn's multiple comparisons, *p<0.05, **p<0.01).

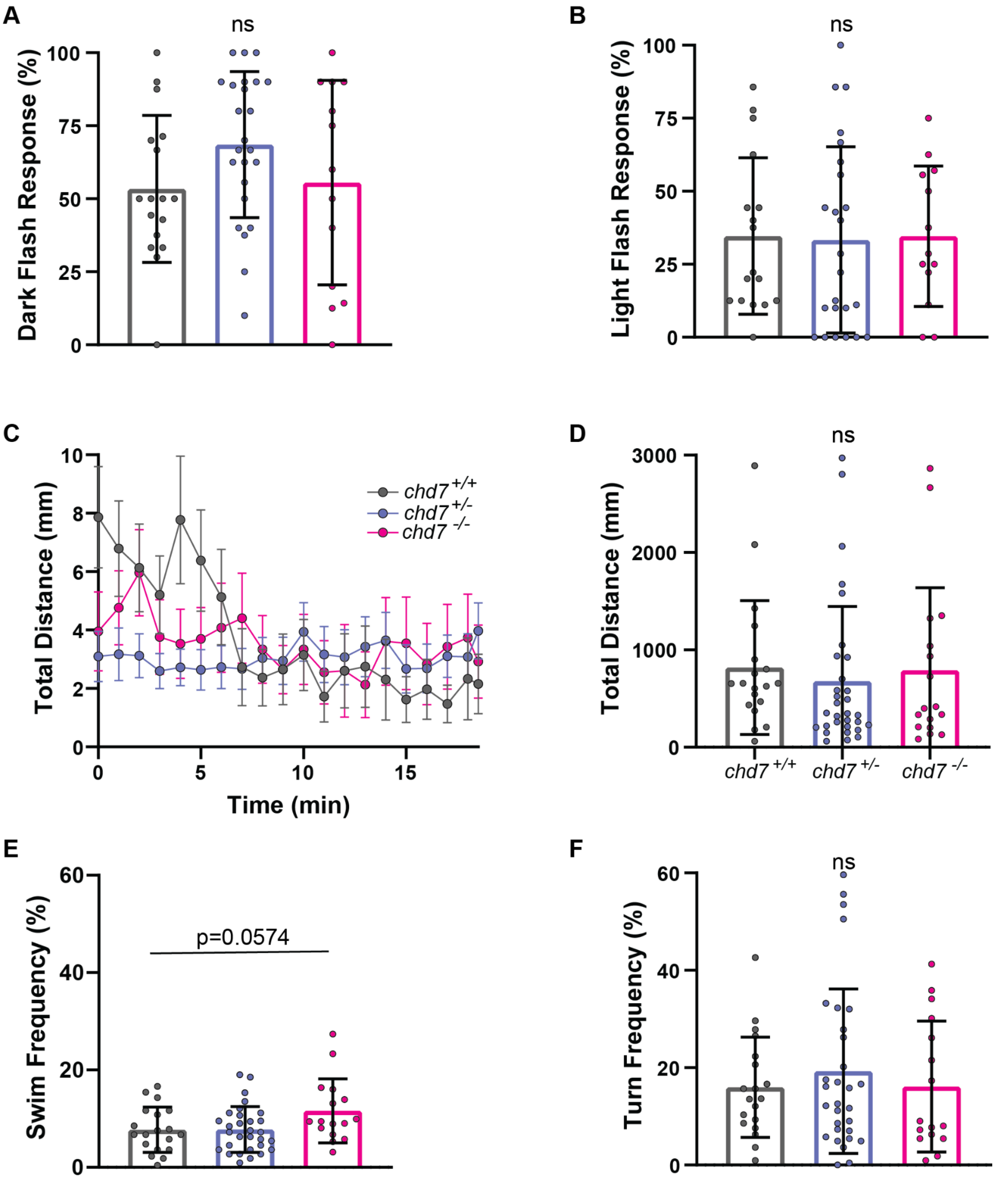
A
ns
Dark Flash Response (%)
B
ns
Light Flash Response (%)
C
Total Distance (mm)
Time (min)
chd7 +/+
chd7 +/-
chd7 -/-
D
ns
Total Distance (mm)
chd7 +/+
chd7 +/-
chd7 -/-
E
p=0.0574
Swim Frequency (%)
F
ns
Turn Frequency (%)

**Supplemental Figure 7. Hyperactivity and visual phenotypes in Exon 16 mutants are independent of strain.** **(A)** Dark flash and **(B)** Light flash response frequencies of Exon 16 mutants propagated in the TLF background (*chd7*$^{+/+}$ n=17, *chd7*$^{+/-}$ n=24, *chd7*$^{-/-}$ n=13) (mean ± SD, Ordinary one-way ANOVA with Tukey's multiple comparison) **(C)** Average total distance traveled plot during 18.5 minutes of recording, represents one testing clutch for *chd7*$^{+/+}$*, chd7*$^{+/-}$*,* and *chd7*$^{-/-}$ (mean ± SEM). **(D)** Sum of total distance traveled for individual larvae, **(E)** swim and **(F)** turning frequencies (*chd7*$^{+/+}$ n= 19, *chd7*$^{+/-}$ n=31, *chd7*$^{-/-}$ n=17) (mean ± SD, Ordinary one-way ANOVA with Tukey's multiple comparison).

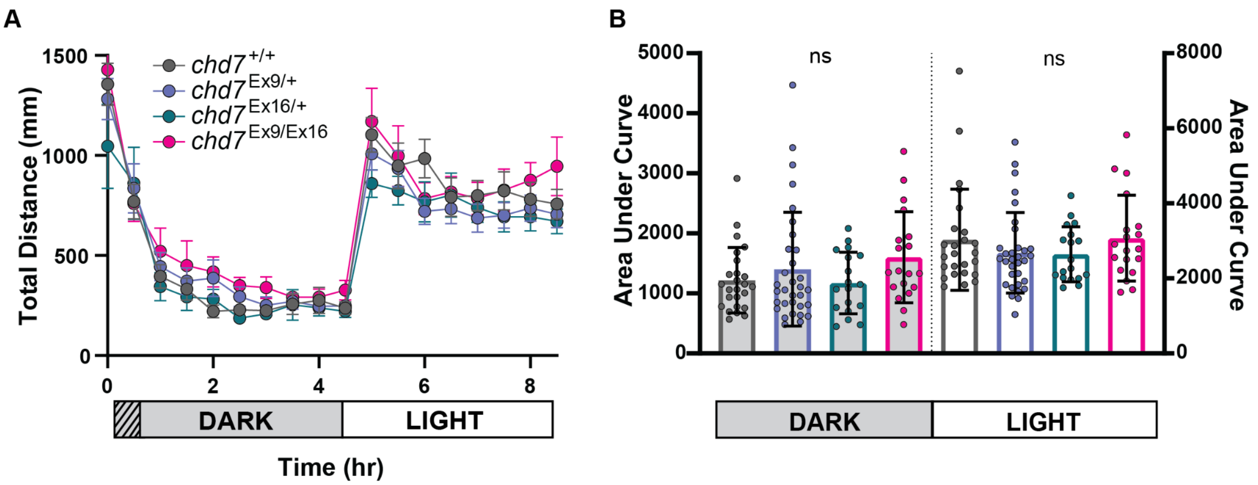

**Supplemental Figure 8. Exon 9/Exon16 double heterozygotes complement Exon 9 hypoactivity.** **(A)** Average total distance traveled plot during 9 hours of recording, analyzed every 30 minutes, including 1 hour of acclimation, 4 hours in the dark, and 4 hours in light (*chd7*$^{+/+}$ n= 25, *chd7*$^{ncu101/+}$ n=32, *chd7*$^{rdu1002/+}$ n=18, *chd7* $^{ncu101/rdu1002}$ n=22) (mean ± SEM). **(B)** Area under the curve for individual distance plots during the dark or light phase, (mean ± SD, Ordinary one-way ANOVA with Dunnett's multiple comparisons).

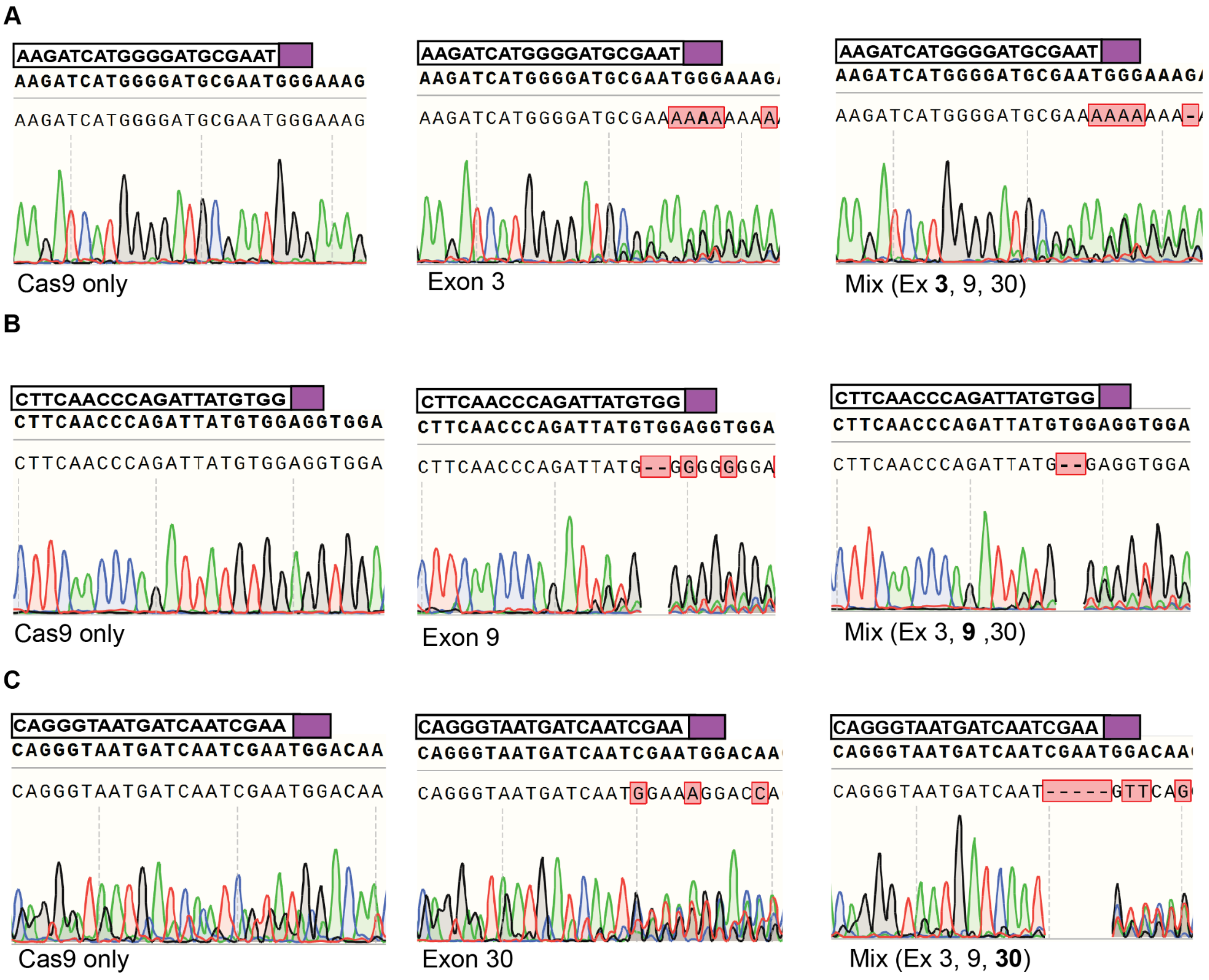

**Supplemental Figure 9. Multiple gRNAs in CRISPANT experiment induce targeted mutagenesis.** Example chromatograms from pooled 5-6 dpf CRISPANTS including **(A)** Cas9 only control injection (Exon 3), Exon 3 gRNA injection, and gRNA mix injection (Exon 3); **(B)** Exon 9 and **(C)** Exon 30; gRNA sequence included with PAM site highlighted with purple box.

| | Craniofacial defect | Otolith defect | Pericardial edema | Noninflated swim bladder | Larvae with any morphological phenotype |
|---|---|---|---|---|---|
| $chd7^{+/-}$ | 1/42, 2.4% | - | 2/42, 4.8% | 3/42, 7.1% | 5/42, 11.9% |
| $chd7^{-/-}$ | 6/19, 31.6% | - | 11/19, 57.9% | 9/19, 47.4% | 15/19, 78.9% |

**Supplemental Table 1. Morphological phenotype frequencies in Exon 16 mutants.**

Morphological phenotype ratios and frequencies in 5-6 dpf $chd7^{rdu1002/-}$ and $chd7^{rdu1002/rdu1002}$; AB background strain.

| | Craniofacial defect | Otolith defect | Pericardial edema | Noninflated swim bladder | Larvae with any morphological phenotype |
|---|---|---|---|---|---|
| $chd7^{Ex9/+}$ | 1/57, 1.8% | 2/57, 3.5% | 2/57, 3.5% | 8/57, 14.0% | 10/57, 17.5% |
| $chd7^{Ex16/+}$ | 1/41, 2.4% | 1/41, 2.4% | 5/41, 12.2% | 5/41, 12.2% | 9/41, 21.9% |
| $chd7^{Ex9/Ex16}$ | 9/42, 21.4% | 5/42, 11.9% | 9/42, 21.4% | 15/42, 35.7% | 25/42, 59.5% |

**Supplemental Table 2. Morphological phenotype frequencies in Exon 9/Exon 16 complementation test larvae.** Morphological phenotype ratios and frequencies in 5-6 dpf $chd7^{ncu101/+}$, $chd7^{rdu1002/+}$, $chd7^{ncu101/rdu1002}$.

| | Application | Target location | Phenotypes | Citation/ Source |
|---|---|---|---|---|
| **ZEBRAFISH** | CRISPR | Exon 9, CHROMO domain 2 | pericardial edema, otolith defects, craniofacial defects, reduced long-latency c-bends, visual and locomotor response deficits | This manuscript |
| | CRISPR | Exon 16, ATP helicase domain | pericardial edema, craniofacial defects, small eyes, increased locomotor activity | This manuscript (gift of Dr. Erica Davis) |
| | CRISPR | Exon 17, ATP helicase domain | small head, 20% pericardial edema, cranial-facial malformations, increased locomotor activity, decreased GABAergic development | Jamadagni, et al. 2021; *EMBO reports* |
| | CRISPR | Exon 2, Low complexity #1 | Pericardial edema, uninflated swim bladder, melanocyte patterning defect, reduced vagal and enteric innervation | Cloney, et al. 2018; *FEBS J* |

| | | | | |
|---|---|---|---|---|
| **MORPHANTS, ZEBRAFISH** | Splice blocking morpholino | Exon 8, Intron 8, CHROMO domain 1 | craniofacial defects, pericardial edema, pigmentation defects, impaired neural crest cell migration | Asad, et al. 2016; *Human Molecular Genetics* |
| | Splice blocking morpholino | Exon 13, Intron 13, ATP helicase domain | T-cell defect, coloboma, otolith defects, cartilage defects, cardiac defects | Liu, et al. 2018; *American Journal of Pathology* |
| | Translation and splice blocking morpholinos | Exon 2, UTR;<br>Exon 13, Intron 13, ATP helicase domain | flattening of head, abnormal tail fins, smaller eyes and disrupted retinal organization, pericardial edema, otolith defects | Patten, et al. 2012; *PLoS One* |
| | Splice blocking morpholino | Intron 15, Exon 16, ATP helicase domain | pectoral fin defects, eye abnormalities, otolith defects, pericardial edema | Balow, et al. 2013; *Developmental Biology* |
| **MOUSE** | Conditional Knockout | Exon 3, Low complexity #10 | motor delay, coordination deficits, cerebellar hypoplasia and foliation anomalies | Whittaker, et al. 2017; *Journal of Clinical Investigation* |
| | ENU mutagenesis | Exon 11, ATP helicase domain | inner ear malformations, slowed growth, eye abnormalities, genital hypoplasia, cardiovascular abnormalities, cleft palate, choanal defects, circling behavior | Bosman, et al. 2005; *Human Molecular Genetics* |
| | ENU mutagenesis | Exon 11, ATP helicase domain | decreased olfactory bulb length, hypogonadotropic hypogonadism, reproductive organ defects, anosmia | Bergman, et al. 2010; *European Journal of Human Genetics* |
| **XENOPUS** | Translation blocking morpholino | chr6L:117963100..117963124 | eye coloboma, otolith defects, craniofacial defects, heart defects, improper neural crest cell migration | Bajpai & Wysocka, et al. 2010; *Nature* |
| **C. ELEGANS** | RNAi, KO | Multiple targets | undetached cuticle, gonad migration defects, stunted length | Jofre, et al. 2022; *PLoS One* |
| **DROSOPHILA** | RNAi | Exon 3, multiple targets, Low complexity #9 | wing abnormalities, defective climbing behavior, memory deficiency, improper axon pruning/migration, defective photoreceptor axon migration, eye defects, mislocalized dorsal cluster neurons | Melicharek, et al. 2010; *Human Molecular Genetics* |
| | EMS Mutagenesis | 5’ end of gene (allele 1), Exon 3 (allele 2), Low complexity #9 | presynaptic motor neuron growth, increase in cell adhesion proteins and postsynaptic proteins, decreased localization of glutamate receptors, stunted synaptic transmission, misaligned presynaptic and | |

| | | | |
|---|---|---|---|
| | | | postsynaptic neurons, decreased movement |

**Supplemental Table 3. Comparison of phenotypes and mutations across CHARGE syndrome models.** Morphological phenotypes, genetic editing application, and mutation locations exhibited in different *chd7* mutant animal models. Represented animal models include zebrafish, *xenopus,* mouse, *c. elegans,* and drosophila. Mutagenesis techniques include N-ethyl-N-nitrosourea (ENU) application, ethyl methanesulfonate (EMS) application, RNA interference (RNAi), morpholinos, and CRISPR/Cas9.

| **Targeted Exon** | **Ensembl GRCz11 Locus** | **Sequence (5'-3')** | **Protein binding domain (SMRT)** |
|---|---|---|---|
| Exon 3 | 2:21785410-21785432(-) | AAGATCATGGGGATGCGAAT**GGG** | Low complexity 14 |
| Exon 9 | 2: 21783574-21783596(-) | CTTCAACCCAGATTATGTGG**AGG** | CHROMO domain 2 |
| Exon 30 | 2:21769382-21769404(-) | CAGGGTAATGATCAATCGTA**TGG** | Low complexity 24 |

**Supplemental Table 4.** Designed gRNA sequences including exon, locus, strand, sequence with PAM site bolded, and corresponding domain.

| **Gene** | **Forward primer (5'-3')** | **Reverse primer (5'-3')** | **Amplicon size(s)** |
|---|---|---|---|
| Exon 9 sequencing | GAGAGGTATATGCCACCCACAT | CAGATGGTTTGAGAACGATTGA | 289 bp |
| Exon 3 sequencing | TCTAGTCGGCAGGTGAAGAGG | CTGGGGCAGGATTTGAGACAG | 465 bp |
| Exon 30 sequencing | GAGCTTTGAGCAAACCAAGGA | GGACTTGCCTTGTTGACCTG | 451 bp |
| 7bp deletion genotyping | GATGATGAGCCCTTCAACCCAG | CAGATGGTTTGAGAACGATTGA | WT 132 bp<br>HET 132/ 125 bp<br>MUT 125 bp |
| 1bp deletion genotyping * | GCAAGTGGCACAAATGTAAAAA | TAAGGATCCTACCGTCTC TGGA | WT 349 bp<br>HET 349/ 254 bp<br>MUT 254 bp |
| *Chd7* cDNA | CAGATGGTGTCGAGGGAGAGG | AGATTGCCTGGCTTCCCGTT | 172 bp |
| *Eif1a* cDNA | CTGGAGGCCAGCTCAAACAT | TCAAGAAGAGTAGTACCGCTAG CATTAC | 86 bp |

**Supplemental Table 5.** Primers for PCR and qPCR including amplicon sizes. *For 1 base pair deletion detection, additional enzyme digest using *MfeI* was required to resolve bands. 1bp deletion created unique *MfeI* restriction site.

| Sequence | PAM | #MM | Gene | Locus | Forward primer | Reverse primer |
|---|---|---|---|---|---|---|
| TTTCAATCCAGATTATG TGG | AAG | 2 | *chd8* | 2:-3811128 0 | TATGTGGTGTGTC TGCTTCAGG | GTGGTTGATAAATG CCTACCTCC |
| CTATAATCCAGATTATA TGG | CAG | 4 | *plk4* | 17:+434136 78 | ACTTATTGTGCCC AAATGGCT | TGATACCGAGAAAA TCTGTGTTGA |
| TTTCAATCCAGATTATA TTG | AGG | 4 | *chd6* | 6:-5162555 5 | TGGGTCAATTTT GGCCCGTA | GCTCCCAAGTGCTC TCTTCA |

**Supplemental Table 6.** Predicted off-target sites using CHOPCHOP from exon 9 gRNA including PAM sites, off-target score, number of mismatches, gene, locus, and sequencing primers. *chd8* and *chd6* targets are within an exon, *plk4* target is an intron.
